# Hippocampal subfield volumes across the healthy lifespan and the effects of MR sequence on estimates

**DOI:** 10.1101/2020.05.28.121343

**Authors:** Aurelie Bussy, Eric Plitman, Raihaan Patel, Stephanie Tullo, Alyssa Salaciak, Saashi A. Bedford, Sarah Farzin, Marie-Lise Béland, Vanessa Valiquette, Christina Kazazian, Christine Tardif, Gabriel A. Devenyi, Mallar Chakravarty

**Affiliations:** Computional Brain Anatomy (CoBrA) Laboratory, Cerebral Imaging Centre, Douglas Mental Health University Institute, Montreal, Quebec, Canada; Integrated Program in Neuroscience, McGill University, Montreal, Canada; Department of Psychiatry, McGill University, Montreal, Quebec, Canada; Department of Biomedical Engineering, McGill University, Montreal, Quebec, Canada; McConnell Brain Imaging Centre, Montreal Neurological Institute, Montreal, Quebec, Canada; Department of Neurology and Neurosurgery, McGill University, Montreal, Quebec, Canada

## Abstract

The hippocampus has been extensively studied in various neuropsychiatric disorders throughout the lifespan. However, inconsistent results have been reported with respect to which subfield volumes are most related to age. Here, we investigate whether these discrepancies may be explained by experimental design differences that exist between studies. Multiple datasets were used to collect 1690 magnetic resonance scans from healthy individuals aged 18-95 years old. Standard T1-weighted (T1w; MPRAGE sequence, 1 mm^3^ voxels), high-resolution T2-weighted (T2w; SPACE sequence, 0.64 mm^3^ voxels) and slab T2-weighted (Slab; 2D turbo spin echo, 0.4 x 0.4 x 2 mm^3^ voxels) images were acquired. The MAGeT Brain algorithm was used for segmentation of the hippocampal grey matter (GM) subfields and peri-hippocampal white matter (WM) subregions. Linear mixed-effect models and Akaike information criterion were used to examine linear, second or third order natural splines relationship between hippocampal volumes and age. We demonstrated that stratum radiatum/lacunosum/moleculare and fornix subregions expressed the highest relative volumetric decrease, while the cornus ammonis 1 presented a relative volumetric preservation of its volume with age. We also found that volumes extracted from slab images were often underestimated and demonstrated different age-related relationships compared to volumes extracted from T1w and T2w images. The current work suggests that although T1w, T2w and slab derived subfield volumetric outputs are largely homologous, modality choice plays a meaningful role in the volumetric estimation of the hippocampal subfields.

## 1. Introduction

Medial temporal lobe structures, particularly the hippocampus, have been extensively studied for their involvement in various neuropsychiatric disorders such as Alzheimer’s disease (Pol et al. 2006; Zhao et al. 2019), schizophrenia (Nelson et al. 1997; Heckers 2001), major depression disorder (Campbell and MacQueen 2004; Malykhin et al. 2010), and frontotemporal dementia (Laakso et al. 2000; Muñoz-Ruiz et al. 2012). To better contextualize group differences in case-control studies, an understanding of normative variation of hippocampal structure across the adult lifespan is critical. This information is crucial in order to better understand how deviations from this trajectory may precede the frank onset (or even the prodromes) of various neuropsychiatric disorders. Unfortunately, current studies investigating the relationship between hippocampus structure and age have rendered inconsistent results.

An examination of studies that report upon the relationship between hippocampal volume and age in healthy aging highlights these inconsistencies: some studies report global preservation (Sullivan et al. 1995; Sullivan et al. 2005; Good et al. 2001), while others report overall reduction (Raz et al. 2004; Kurth et al. 2017; Malykhin et al. 2017; Bussy et al. 2019). More recently, several studies have begun to characterize this relationship at the level of the grey matter (GM) hippocampal subfields, including cornu ammonis (CA)1, 2, 3 and 4, subiculum and stratum radiatum/lacunosum/moleculare (SRLM), using innovative new techniques (Goubran et al. 2013; Winterburn et al. 2013; Pipitone et al. 2014; Yushkevich et al. 2015; Olsen et al. 2019). In most studies reviewed in de Flores et al. (2015), the CA1 and subiculum appear to be most impacted by aging. However, in other recent studies, the CA1 (Voineskos et al. 2015; Amaral et al. 2018) and subiculum (Daugherty et al. 2016) have been shown to be relatively preserved. In the present work, we suggest that the observed discrepancies in findings may be explained by the study design disparities - across a range of methodological choices - that exist between studies.

First, a key methodological difference is the anatomical definition of the hippocampal subfields. This issue is controversial and thus, different atlases are used in the literature (Winterburn et al. 2015; Iglesias et al. 2015; Yushkevich et al. 2015; Yushkevich et al. 2015; Amaral et al. 2018; Palombo et al. 2013). In order to facilitate the segmentation of small subfields, protocols use diverse definitions of subfields, which include CA1 combined with CA2 (Bender et al. 2018), CA3 combined with dentate gyrus (DG) (de Flores et al. 2015), CA3 combined with CA4 and DG (Shing et al. 2011) or CA4 solely combined with DG (Wisse et al. 2012; Winterburn et al. 2015). Second, various segmentation protocols are employed to study the subfields, including manual delineation (Mueller et al. 2007; La Joie et al. 2010), semi-automated approaches (Yushkevich et al. 2010), and automated methods (Van Leemput et al. 2009; Pipitone et al. 2014; Iglesias et al. 2015; Yushkevich et al. 2015).

Of note, a recent effort has been started by the Hippocampal Subfields Group (http://www.hippocampalsubfields.com/) to develop a harmonized protocol based on expert consensus and histological evidence as a means to facilitate comparison of findings across subject groups (Wisse et al. 2017; Olsen et al. 2019). However, despite these significant and important efforts, there remains two additional aspects related to study design that are outside the scope of segmentation protocols, namely: 1) the non-linear relationship of the hippocampal subfield volumes with age, 2) the magnetic resonance image (MRI) acquisition protocols. Further, to completely characterize the age-related trends of the hippocampal circuitry, it is critical to examine the examination of peri-hippocampal white matter (WM) subregions such as alveus, fornix, fimbria and mammillary bodies (MB). The current manuscript primarily focuses on the impact of these two components.

Importantly, previous studies have examined diverse participant age ranges. While some cohorts consider the entire adult lifespan (Mueller et al. 2007; de Flores et al. 2015; Amaral et al. 2018; Zheng et al. 2018), others only assess subjects older than 65 (Frisoni et al. 2008; Wisse et al. 2014). These variations can lead to inconsistencies in findings, since it is well-accepted that the relationship between brain structures and age is typically non-linear (Coupé et al. 2017; Tullo et al. 2019); thus, sampling a smaller age range can lead to conclusions that should not be used to interpret data outside that range. Nonetheless, only a few studies have investigated the non-linear relationships between age and hippocampal subfield volumes (Mueller et al. 2007; Ziegler et al. 2012; de Flores et al. 2015; Malykhin et al. 2017).

MRI protocols are highly heterogeneous in the literature and its impact on hippocampal subfield definitions is still not clear. Indeed, studies have used standard T1-weighted (T1w) images (Frisoni et al. 2008; Chételat et al. 2008; Amaral et al. 2018), more specialized T2-weighted (T2w) images (Mueller et al. 2007; Wisse et al. 2014) or proton-density-weighted (PDw) images (La Joie et al. 2010; de Flores et al. 2015). Further, the resolution used to examine hippocampal subfields is highly variable across these image acquisitions. For example, studies used isotropic whole brain scans at 0.7 mm^3^, 1 mm^3^ (Wisse et al. 2014; Pereira et al. 2014), (typically for T1w images, and rarely T2w) or 0.78 x 0.78 x 1.5 mm^3^ resolution (Voineskos et al. 2015), or “slab” scans at 0.4 x 0.4 x 2 mm^3^ resolution (Mueller and Weiner 2009) (typically for T2w and PD images). The latter is acquired in a region of interest that covers the amygdala and hippocampus either partially or entirely from its anterior to posterior extent. Taken together, these different acquisition parameters may contribute to volume estimation differences potentially explaining the variation observed in age-related subfield relationships. Further, there are very few studies that used high-resolution isotropic scans which allow for the visualization of the molecular layers (i.e. “dark band”) of the hippocampus, often considered a prerequisite landmark for manual or automated identification of hippocampal subfields (Eriksson et al. 2008; Goubran et al. 2014; Winterburn et al. 2013; Wisse et al. 2017). While there have been suggestions in several studies that slab T2w or PDw scans should be preferentially used in the study of hippocampal subfields to obtain coronal high resolution (La Joie et al. 2010; Yushkevich et al. 2015), it is still unclear whether the slice thickness biased volume measurements. Concurrently, there has also been minimal investigation with respect to the use of higher resolution isotropic images for this purpose (Wisse et al. 2012; Wisse et al. 2014). Finally, given that the hippocampus and its subfields make up a critical circuit in the brain’s memory network, some groups (Iglesias et al. 2015), including ours (Amaral et al. 2018; Tardif et al. 2018), have begun to include the peri-hippocampal WM in studies examining the hippocampal subfields. This is a critical step forward towards examining this specific anatomy at the circuit level.

Given the inconsistency in the literature, there is a clear need to better characterize age-related relationships between the volume of the hippocampal subfields and peri-hippocampal WM subregions while performing a critical assessment of experimental design choices. Here, we examine the role of image acquisition (including standard isotropic T1w, high-resolution isotropic T2w and slab hippocampus-specific T2w) on hippocampal subfield volumes, while assessing the reliability of the hippocampal subfield measures.

## 2. Methods

### 2.1. Image acquisition and participants

#### 2.1.1 Image acquisition types

Here, we examine three different scan types from multiple sources. These data come from multiple datasets collected by our group and publicly available datasets (described further in section 2.1.2). These scan types were targeted because two of them (standard T1w MPRAGE and slab T2-weighted 2D turbo spin echo [TSE]) have been commonly used in the hippocampal subfield literature. The final one, a high-resolution whole brain T2w acquisition, is introduced here as a methodology that potentially addresses some of the limitations with respect to resolution and field-of-view.

- *T1-weighted:* This is the protocol most commonly used in neuroimaging studies and has been extensively employed to study the hippocampus, among other structures (Pereira et al. 2014). Here, we used the Alzheimer’s Disease Neuroimaging Initiative (ADNI) magnetization prepared - rapid gradient echo (MPRAGE) protocol (Mugler and Brookeman 1990; Jack et al. 2008), since these parameters have been commonly used to investigate hippocampus physiology and pathology.
- *Slab T2-weighted 2D TSE:* This protocol uses the same ADNI oblique acquisition with 2 mm thick slices perpendicular to the long axis of the hippocampus and 0.4 mm x 0.4 mm in the coronal plane. Recently, this sequence became the standard procedure to study the hippocampus subfields. Users typically take advantage of the high-resolution of the coronal plane to segment the subfields while sacrificing resolution through the anterior-posterior direction. It is unclear what this design trade-off does in terms of consistency and precision of the measurement.
- *High-resolution 0.64 mm^3^ T2w:* This protocol uses the SPACE sequence, a 3D TSE sequence with slab selective, variable excitation pulse. It has been developed in our laboratory for the purpose of increasing the resolution and subfields contrast in the hippocampus. Furthermore, this sequence provides isotropic images across the whole brain and allows for the use of standard linear and nonlinear registration methods that expect whole-brain coverage (Klein et al. 2009; Chakravarty et al. 2013).

#### 2.1.2 Datasets

The five datasets used to examine the relationship of the hippocampal subfields with age are outlined below, and all include some variation of the acquisition methods described in the previous section. All the datasets collected by our group at the Douglas Mental Health University Institute were approved by its Research Ethics Board.

- *Healthy Aging (HA).* This dataset was collected and scanned on a 3T Siemens Trio MRI scanner using a 32-channel head coil at the Douglas Mental Health University Institute, Montreal, Quebec, Canada, and contains 111 participants aged 18-80 (Tullo et al. 2019). We analyzed two types of MRI images: standard MPRAGE (1 mm^3^) and high-resolution T2-weighted (TSE; 0.64 mm^3^).
- *Alzheimer’s Disease Biomarkers (ADB).* This dataset was also collected and scanned on a 3T Siemens Trio MRI scanner using a 32-channel head coil at the Douglas Mental Health University Institute, Montreal, Quebec, Canada, (Tullo et al. 2019). From this study, we used 68 healthy elderly participants (56-81). The same acquisition protocol as the HA dataset was used, providing standard MPRAGE (1 mm^3^) and high-resolution T2-weighted (TSE sequence; 0.64 mm^3^).
- *Alzheimer’s Disease Neuroimaging Initiative (ADNI).* ADNI is a publicly available multicenter study from which we included 317 healthy participants, aged 56-95, scanned on 3T General Electric, Philips or Siemens scanners, depending on the acquisition’s site. We used two types of MRI images: standard MPRAGE (1 mm^3^) (Jack et al. 2008) and slab T2-weighted 2D TSE with 2 mm thick slices perpendicular to the long axis of the hippocampus and 0.4 mm dimensions in the coronal plane (Thomas et al. 2004; Mueller et al. 2007; Mueller et al. 2018).
- *Cambridge Centre for Ageing and Neuroscience (Cam-CAN).* Cam-CAN is a large-scale dataset collecting MRI scans at the Medical Research Council Cognition and Brain Sciences Unit in Cambridge, England, using a 3T Siemens TIM Trio scanner with a 32-channel head coil (Shafto et al. 2014; Taylor et al. 2017). We included 652 healthy individuals (aged 18-88) with standard MPRAGE (1 mm^3^).
- *Test-retest*. Test-retest dataset was collected at the Douglas Mental Health University Institute, Montreal, Canada. Twenty-one healthy participants aged 20-42 were recruited and scanned using a 3T Siemens PRISMA scanner. These participants underwent three different MR sequence acquisitions that reproduce the three different types of acquisitions described above: standard MPRAGE (1 mm^3^), high-resolution T2-weighted (TSE sequence; 0.64 mm^3^) and slab T2-weighted 2D TSE sequence (0.4 x 0.4 x 2 mm).

Initial and final sample characteristics including number of scans by dataset, age, sex, and sequence are summarized in Table 1. The total number of scans after quality control (QC) of the images (see *Section 3 Image processing),* was 930, and consisted of individuals aged 18-93 (58.4% female) (Table 1B). Complete demographics and QC inclusion/exclusion criteria are available in Supplementary material; age distribution of the participants included in the analyses can be found in Supplementary figure 1. The main acquisition parameters are provided in Table 2 and more details given in Supplementary material - Acquisition parameters.

**Table 1:** Demographics by dataset: **A**) Initial sample, **B**) Final sample after motion and segmentation quality control (QC) across *Healthy Aging (HA), Alzheimer’s Disease Biomarkers (ADB), Alzheimer’s Disease NeuroImaging Initiative (ADNI), and Cambridge Centre for Ageing and Neuroscience (Cam-CAN). **np** = number of participants; **ns** = number of scans.*

| **A** | ***ADB***<br>(*np=68*)<br>(*ns=136*) | ***ADNI***<br>(*np=317*)<br>(*ns=617*) | ***Cam-CAN***<br>(*np=652*)<br>(*ns=652*) | ***HA***<br>(*np=111*)<br>(*ns=222*) | ***Test-retest***<br>(*np=21*)<br>(*ns=63*) | ***Overall***<br>(*np=1169*)<br>(*ns=1690*) |
| --- | --- | --- | --- | --- | --- | --- |
| **Age** |  |  |  |  |  |  |
| *Mean (SD)* | 69.7 (5.7) | 73.9 (8.0) | 54.3 (18.6) | 45.6 (16.4) | 27.0 (5.9) | 60.6 (18.8) |
| *Median* | 70.0 | 72 | 54.5 | 44.0 | 27.0 | 67 |
| *[Min, Max]* | [56.0, 81.0] | [56.0, 95.0] | [18.0, 88.0] | [18.0, 95.0] | [20.0, 42.0] | [18.0, 95.0] |
| **Sex** |  |  |  |  |  |  |
| *F* | 82 (60.3%) | 387 (62.7%) | 330 (50.6%) | 116 (52.3%) | 39 (61.9%) | 954 (58.4%) |
| *M* | 54 (39.7%) | 230 (37.3%) | 322 (49.4%) | 106 (47.7%) | 24 (38.1%) | 736 (41.6%) |
| *Sequence* |  |  |  |  |  |  |
| *T1* | 68 (50.0%) | 317 (51.4%) | 652 (100%) | 111 (50.0%) | 21 (33.0%) | 1169 (69.2%) |
| *T2* | 68 (50.0%) | 0 (0.0%) | 0 (0.0%) | 111 (50.0%) | 21 (33.0%) | 200 (11.8%) |
| *Slab* | 0 (0.0%) | 300 (48.6%) | 0 (0.0%) | 0 (0.0%) | 21 (33.0%) | 321 (19.0%) |

| **B** | *ADB*<br>(*np=52*)<br>(*ns=100*) | *ADNI*<br>(*np=135*)<br>(*ns=233*) | *Cam-CAN*<br>(*np=376*)<br>(*ns=376*) | *HA*<br>(*np=84*)<br>(*ns=167*) | *Test-retest*<br>(*np=18*)<br>(*ns=54*) | *Overall*<br>(*np=665*)<br>(*ns=930*) |
| --- | --- | --- | --- | --- | --- | --- |
| *Age* |  |  |  |  |  |  |
| *Mean (SD)* | 69.3 (5.7) | 71.6 (6.9) | 49.9 (17.2) | 46.4 (16.1) | 26.8 (5.6) | 55.4 (18.5) |
| *Median*<br>*[Min, Max]* | 69.5<br>[56.0, 81.0] | 70.0<br>[56.0, 93.0] | 49.0<br>[18.0, 85.0] | 46.0<br>[18.0, 80.0] | 26.0<br>[20.0, 42.0] | 61.0<br>[18.0, 93.0] |
| *Sex* |  |  |  |  |  |  |
| *F* | 61 (61.0%) | 160 (68.7%) | 196 (52.1%) | 93 (55.7%) | 33 (61.1%) | 543 (58.4%) |
| *M* | 39 (39.0%) | 73 (31.3%) | 180 (47.9%) | 74 (44.3%) | 21 (38.9%) | 387 (41.6%) |
| *Sequence* |  |  |  |  |  |  |
| *T1* | 52 (52.0%) | 135 (57.9%) | 376 (100.0%) | 84 (50.3%) | 18 (33.0%) | 665 (71.5%) |
| *T2* | 48 (48.0%) | 0 (0.0%) | 0 (0.0%) | 83 (49.7%) | 18 (33.0%) | 149 (16.0%) |
| *Slab* | 0 (0.0%) | 98 (42.1%) | 0 (0.0%) | 0 (0.0%) | 18 (33.0%) | 116 (12.5%) |

**Table 2:**
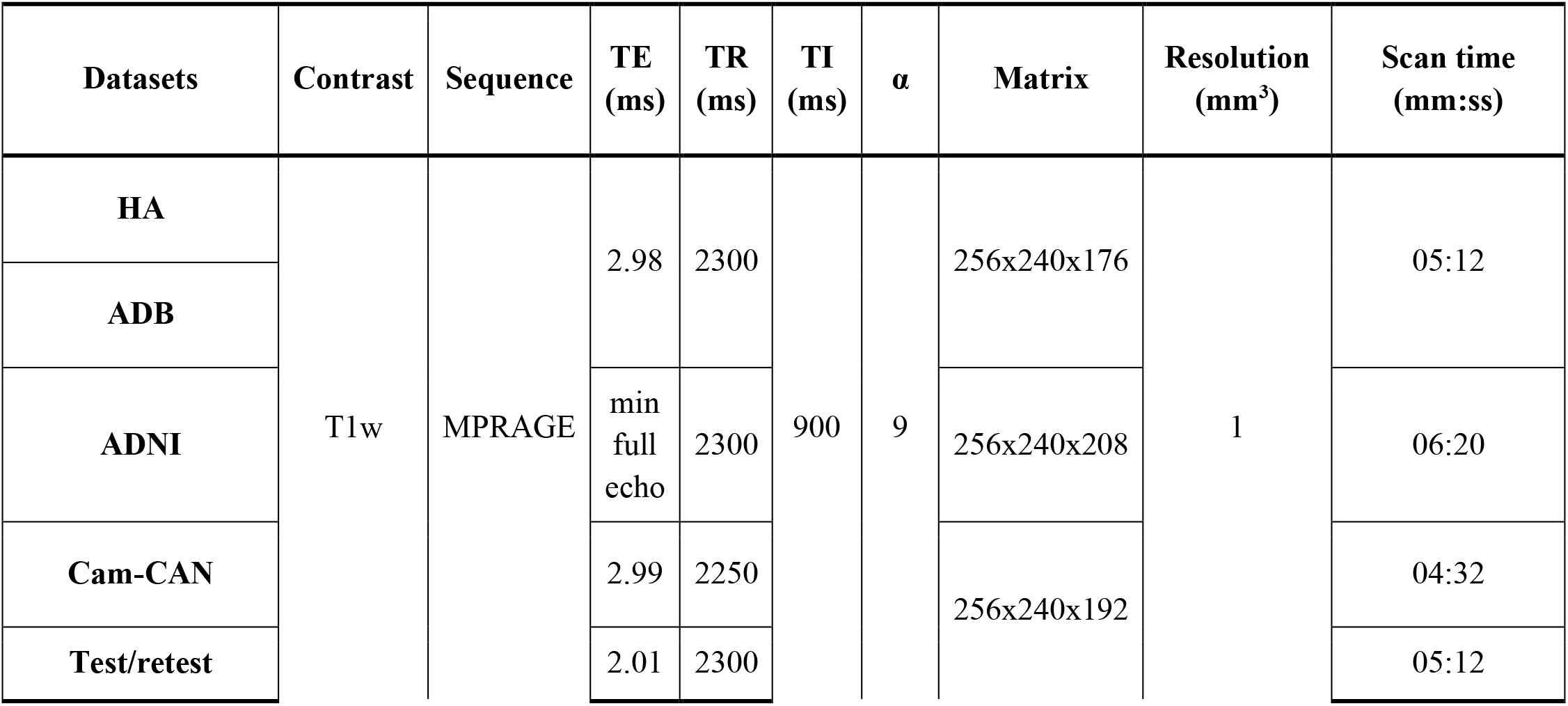
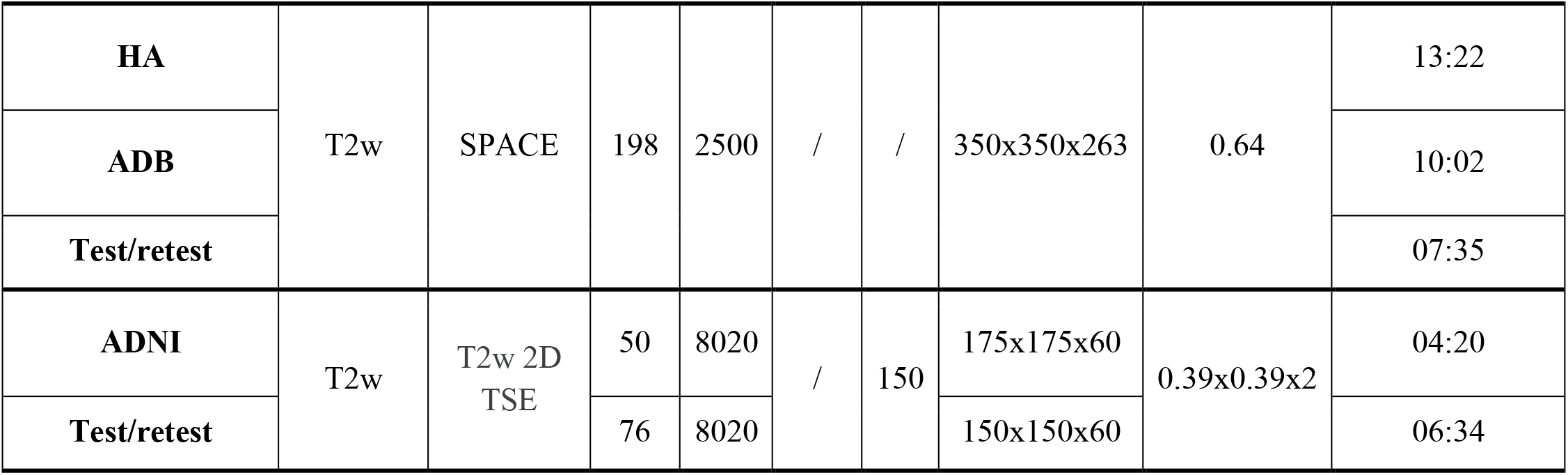
Scanning parameters of the different sequences in the different datasets. TE= echo time, TR= repetition time, TI = inversion time, α= flip angle. More details in Supplementary material - Acquisition parameters.

### 2.2 Image processing

#### 2.2.1 Raw quality control

Structural MR images are particularly sensitive to subject motion, often resulting from involuntary movements (e.g. cardiac or respiratory motion, and drift over time). The effects of motion, including blurring and ringing, negatively impact the quality of structural MRI data (Bellon et al. 1986; Smith and Nayak 2010; Reuter et al. 2015). Quality control (QC) of all raw images was performed by a rater (AB) using the QC procedure previously developed in our laboratory ((Bedford et al. 2020); https://github.com/CoBrALab/documentation/wiki/Motion-Quality-Control-Manual).

#### 2.2.2 Preprocessing

The minc-bpipe-library pipeline (https://github.com/CobraLab/minc-bpipe-library) was employed to preprocess and standardize T1w images using N4 bias field correction (Tustison et al. 2010), registration to MNI space using bestlinreg (Collins et al. 1994; Dadar et al. 2018), standardization of the field-of-view and orientation of the brain using an inverse-affine transformation of a MNI space head mask, and extraction of the brain using BEaST (Eskildsen et al. 2012). The pipeline produces brain masks and quality control images to quickly evaluate all steps of the pipeline.

T2w and slab images preprocessing consists of the following steps: rigid registration of T1w to T2w or slab scan (Collins et al. 1994; Dadar et al. 2018), application of the transform file to the T1w brain masks obtained by the minc-bpipe-library pipeline to create T2w or slab brain masks, N4 correction, and extraction of T2w or slab brains using T2w or slab brain masks.

#### 2.2.3 Automated Hippocampus Segmentation

The Multiple Automatically Generated Templates (MAGeT) Brain algorithm, a modified automated multi-atlas technique (Pipitone et al. 2014; Chakravarty et al. 2013), was used for segmentation of the hippocampal subfields. This method uses a set of five high-quality atlases, manually segmented on 0.3 mm isotropic T1w and T2w brains, as input. We used the Winterburn et al. (2013) definitions of the GM subfields (CA1, combined CA2 and CA3 [CA2CA3], combined CA4 and dentate gyrus [CA4DG], SRLM and subiculum) and the Amaral et al. (2018) definitions of WM subregions (fimbria, fornix, alveus and MB) of the hippocampus (Figure 1).

**Figure 1:**
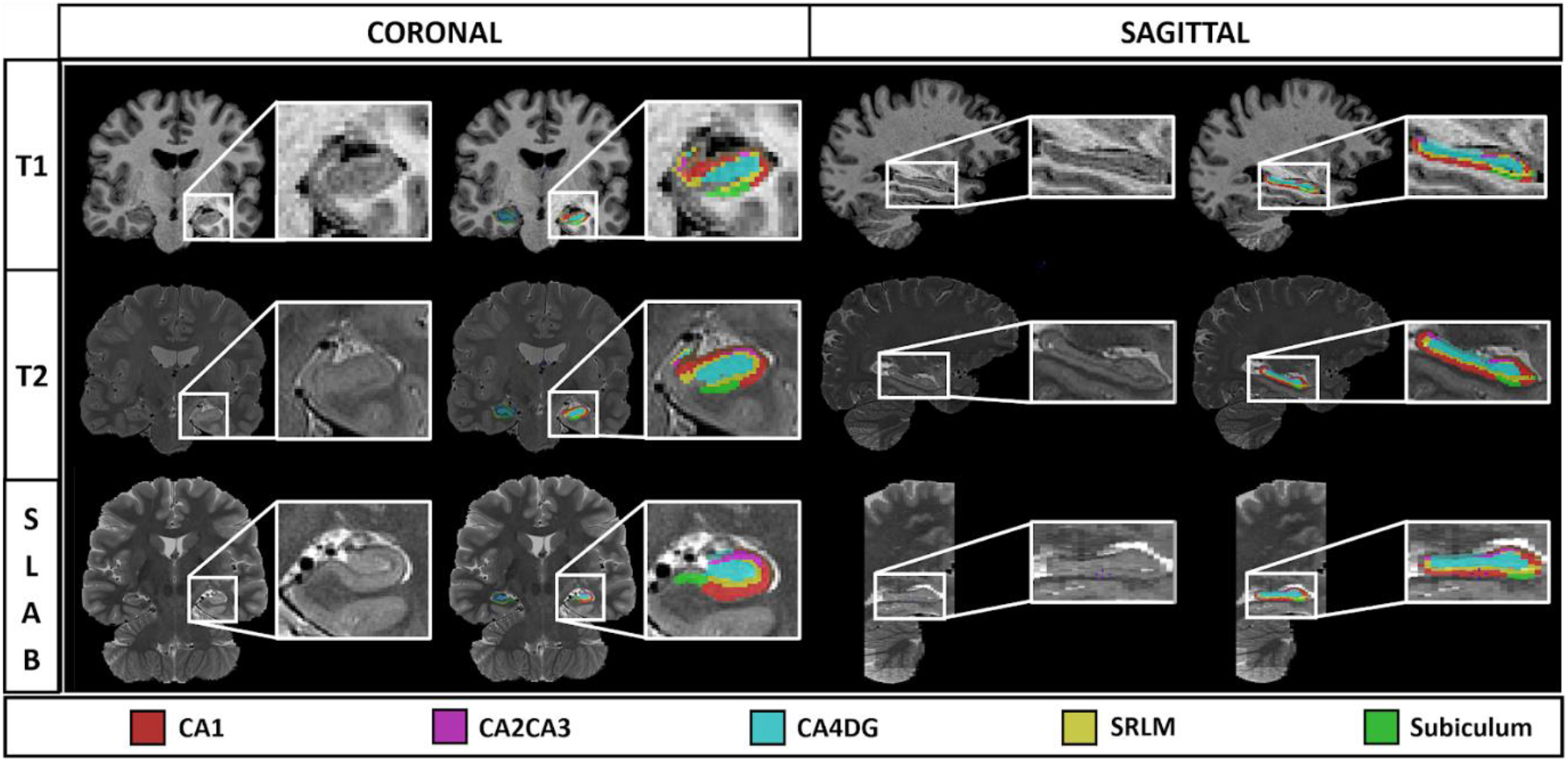
Example of coronal and sagittal views of a participant’s scans: T1w (1 mm^3^), T2 (0.64 mm^3^) and slab (0.4 x 0.4 x 2mm) without and with the labels obtained from our segmentation protocol.

For each sequence-type within a specific dataset (e.g. high-resolution T2w data in the ADB dataset), we first ran a “best template selection” stage (https://github.com/CoBrALab/documentation/wiki/Best-Templates-for-MAGeT), in order to select the 21 subjects with highest quality atlas-to-template segmentation. These 21 subjects were then used as a template library to segment all the subjects. This step artificially increases the number of atlases to 105 (21 templates x 5 atlases) in order to improve segmentation by reducing error propagation compared to traditional atlas-based segmentation procedures (Iosifescu et al. 1997; Svarer et al. 2005). Finally, the 105 candidate labels for each subject were fused using a majority vote technique to obtain final labels (Pipitone et al. 2014; Chakravarty et al. 2013; Makowski et al. 2018). MAGeT Brain uses affine and SyN nonlinear registration, which are registration options provided as part of the Advanced Normalization Tools (ANTS; [Avants et al. 2008]). Segmentations were conducted using a focused region-of-interest (ROI) based registration step for each hemisphere independently. ROI masks were generated by converting all labels to a single value mask and then dilated with a 3 mm radius kernel in order to focus both affine and nonlinear registration, a method which reduces computational time and improves segmentation accuracy (Chakravarty et al. 2008; Chakravarty et al. 2009).

Adaptation of the standard version of MAGeT Brain was performed to process slab scans. We took advantage of the availability of whole brain T1w scans for each of our slab images to provide bulk alignment of the atlases and templates. First, within-subject affine registration was performed between slab and whole brain T1w images by using the label masks to handle mismatched field-of-view. Then, affine registration was performed between subject whole brain T1w images. Finally, nonlinear registrations were computed between the slabs using concatenated affine transforms (within-subject between slab and T1w, and between subject whole brain T1w affine registrations) for initialization.

The rater (AB) quality controlled the final labels by visual inspection following the QC procedure implemented by our group (**https://github.com/CobraLab/documentation/wiki/MAGeT-Brain-Quality-Control-(QC)-Guide**) to only include high quality segmentation in the statistical analyses.

### 2.3 Statistical analysis

#### 2.3.1 Relationship between age and normalized hippocampal subfield volumes

To draw general conclusions on the association between each subfield volume with age, we first combined all the datasets together in order to account for the variability in image resolution, sequence, and age range commonly encountered in the literature (Yushkevich et al. 2015). Linear mixed-effect models (lmer from *lmerTest_3.1-0* package in R 3.6.1) and natural spline (ns from *splines* package) were used to examine the relationship between hippocampal volumes and age. To study different age relationships for each structure of interest, the Akaike information criterion (AIC; Akaike 1974; *stats* package) was used to find the relative quality of each statistical model using either linear, second, or third order natural splines. To minimize the loss of information, the model with the lowest AIC was considered the best fit for the data (Mazerolle 2006).

Sex was used as a fixed effect, and dataset (ADB, ADNI, HA, Cam-CAN, test-retest), MRI sequence (T1, T2, slab), and subject were modelled as random effects for all statistical analyses. In addition, volumetric normalization was performed to account for head size variability across participants. In addition, intracranial volume (ICV) was used to account for interindividual variability in head size (de Flores et al. 2015; Daugherty et al. 2016). Since we initially wanted to investigate the relationship between age with each subfield volume, as well as with total hippocampal GM or WM volume, while covarying for the z-scored ICV (*scale* function) to help the fit of the model (1).

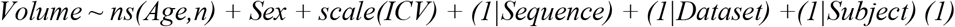

To determine the extent to which each subfield participates in global hippocampus atrophy with age, we then assessed the relationship between each subfield volume and age, while covarying for total hippocampal GM or WM volume as well as ICV (2).

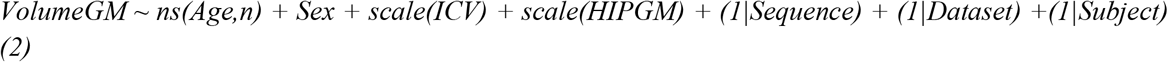

For each analysis described above, we used a Bonferroni correction to correct for multiple comparisons across our 18 structures (five GM subfields and four WM subregions per hemisphere), at a p<0.05 threshold for significance, resulting in a significance level of p<0.0028 (only corrected p-values reported throughout this paper).

To visualize these results, we used two different techniques. The first used these coefficients to create the predicted volume divided by the predicted volume at age 18 for a subject of mean ICV and mean ipsilateral hippocampal GM or WM volume when applicable. This allows us to assess the age relative volume change of each subfield with respect to its baseline volume (Figures 2, 3, 4 and Supplementary figures 6 and 7). The second type of representation was done using the fixed effect coefficients of the statistical models described above to create the corresponding relationship of the volume with age (Supplementary figures 3, 4, 5, 7 and 9). Of note, both visualization techniques show similar shapes with the first relating these slopes in percent volume change with age whereas the second illustrates volume change with age.

**Figure 2:**
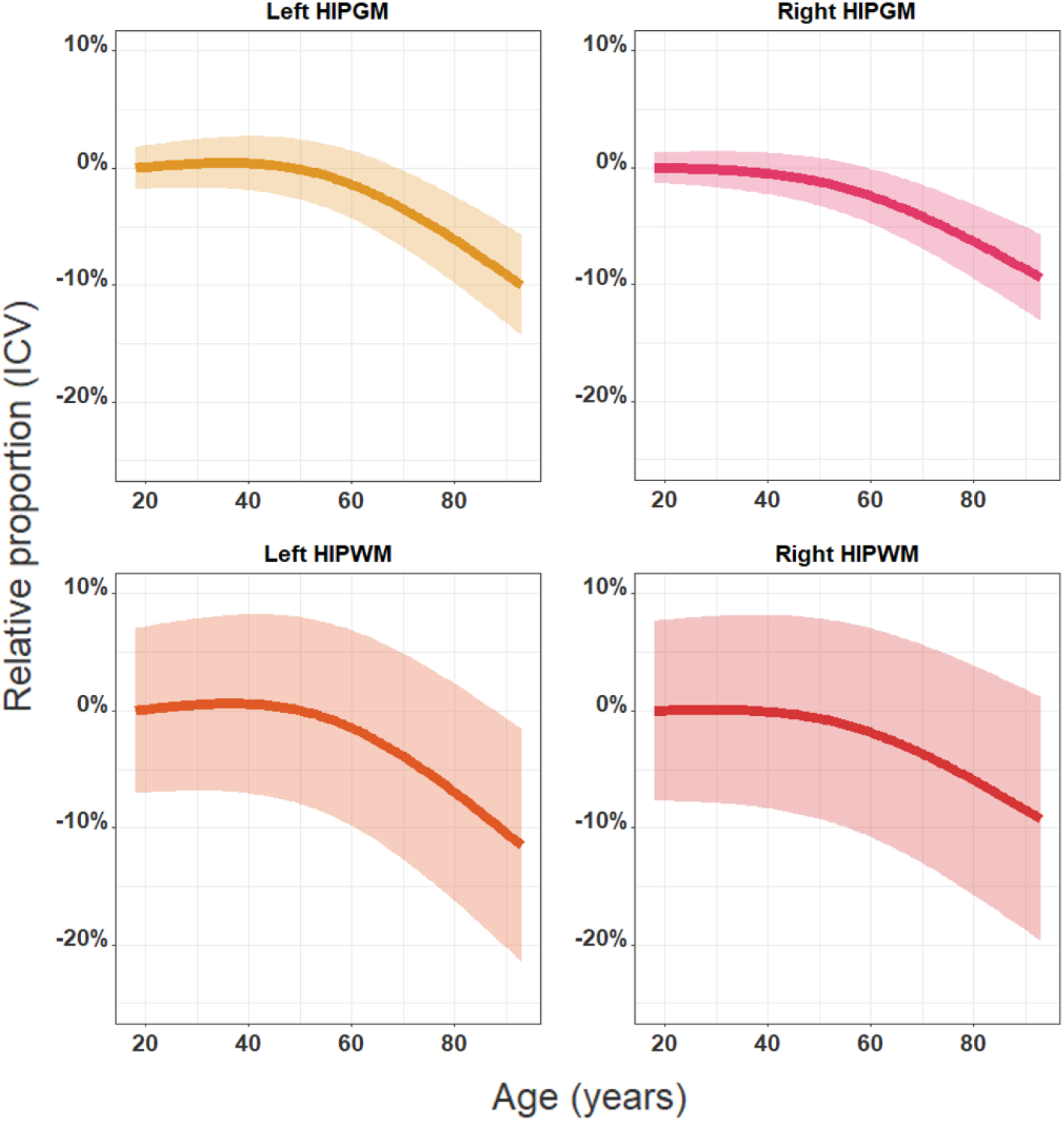
Best fit models showing the relationships between age and the relative proportion of the hippocampus using the predicted volume at age 18 for a subject of mean ICV as baseline (same model with volume instead of relative proportion in Supplementary figure 3). Best fit models displayed for each structure covaried by ICV and sex as fixed effects and dataset, sequence, and subjects as random effects. Second order relationships were found to be the best fit model for all the structures: right HIPGM (p=3.33×10^−4^), right HIPWM (p=1.05×10−^3^), left HIPGM (p=2.27×10−^5^) and left HIPWM (p=1.18×10−^5^).

**Figure 3:**
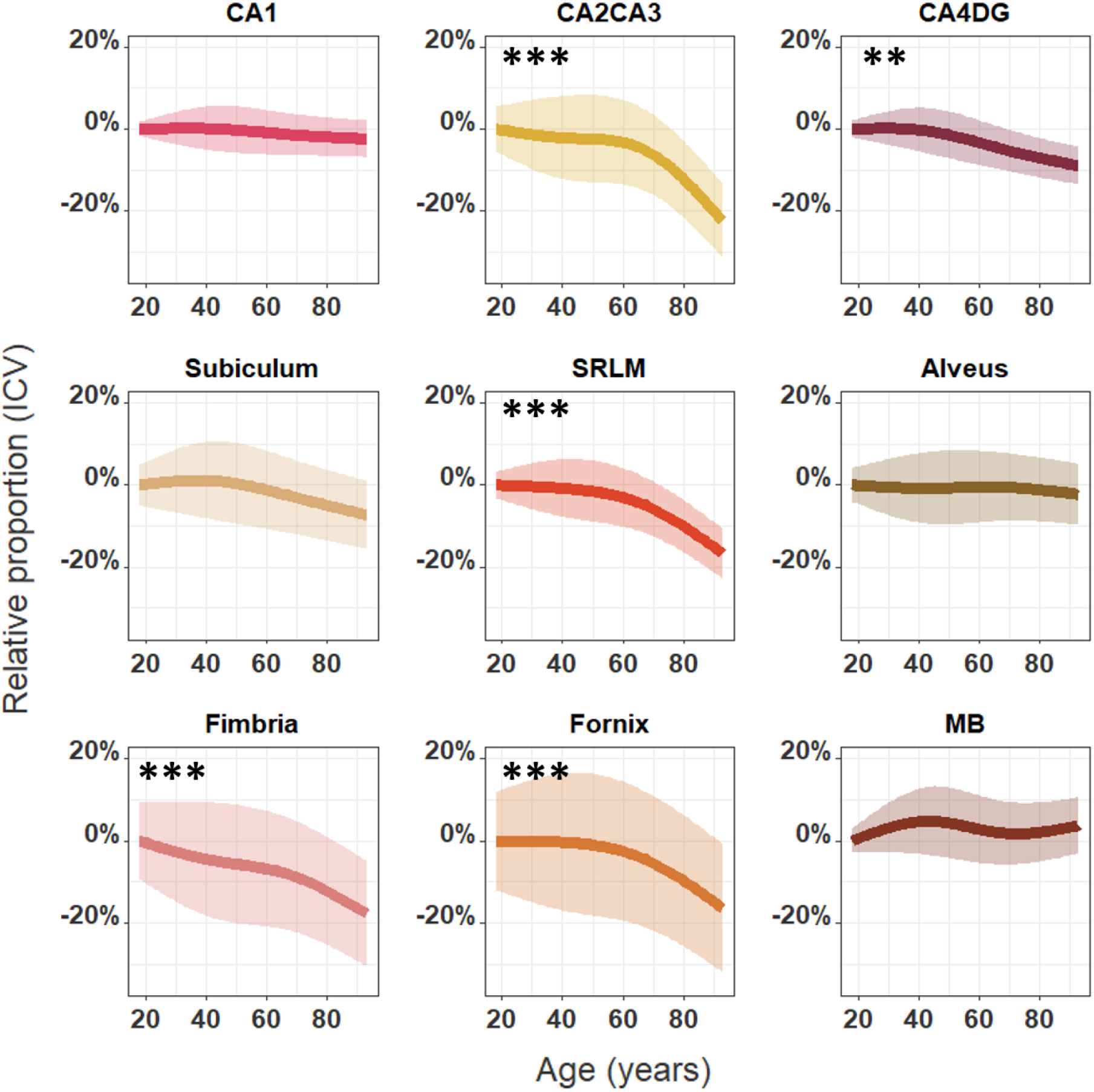
Best fit models showing the relationships between age and the relative proportion of the right hippocampal subfields, using the predicted volumes at age 18 for a subject of mean ICV as baseline (same model with volume instead of relative proportion in Supplementary figure 4). Best fit models displayed for each subfield covaried by ICV and sex as fixed effects and dataset, sequence, and subject as random effects. Significant monotonic decreases were found for the right CA2CA3 (p=3.55×10^−7^), CA4DG (p=4.28×10^−3^), SRLM (p=7.74×10^−7^), fimbria (p=9.45×10^−4^) and fornix (p=1.79×10^−4^). Similar relationships were found in the left hemisphere (Supplementary figures 6 and 7) * p<0.05; ** p<0.01 and *** p<0.001 after Bonferroni correction.

**Figure 4:**
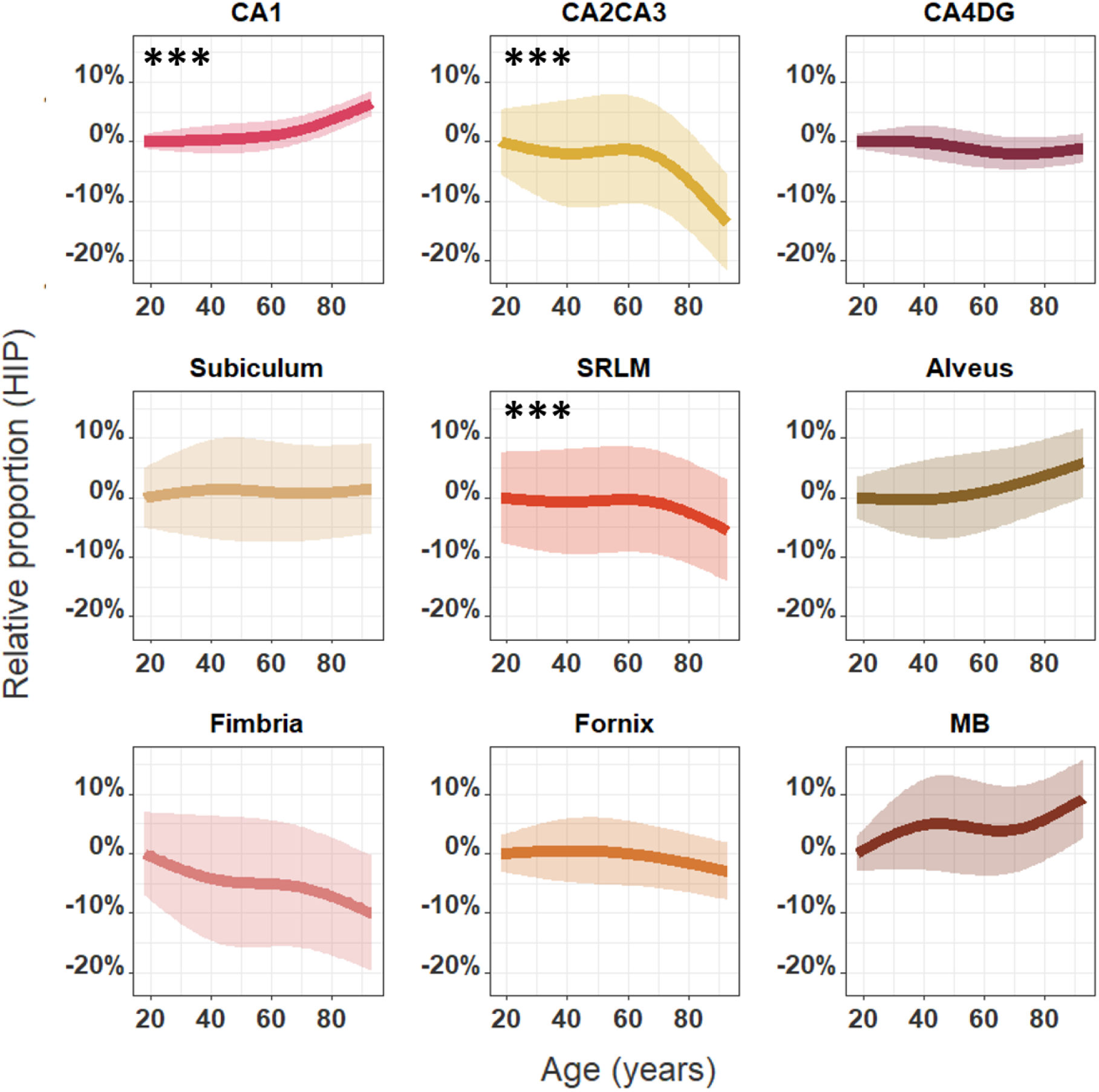
Best fit models showing the relationships between age and the relative proportion of the right hippocampal subfields, using the predicted volumes at age 18 for a subject of mean ICV and mean right hippocampal GM or WM volume as baseline (same model with volume instead of relative proportion in Supplementary figure 5). Best fit model displayed for each subfield covaried by right hippocampal GM or WM volume, ICV and sex as fixed effects and dataset, sequence, and subject as random effects. Significant monotonic increases were found for the right CA1 (p=1.19×10^−10^) and significant monotonic decreases were found for the right CA2CA3 (p=3.19×10^−4^) and SRLM (p=5.29×10^−6^). Similar investigations made in the left hemisphere can be found in Supplementary figures 8 and 9. * p<0.05; ** p<0.01 and *** p<0.001 after Bonferroni correction.

Apolipoprotein E ε4 allele (*APOE4*) gene is often studied in aging population since it is associated with higher risk of both early-onset and late-onset sporadic AD (Corder et al. 1993) as well as age-related cognitive impairment (Rawle et al. 2018). While some authors described that *APOE4* allele was associated with hippocampal, amygdalar and entorhinal cortex atrophy (Cherbuin et al. 2007), others did not find any structural differences between *APOE4* carriers and noncarriers (Habes et al. 2016) or even that *APOE4* noncarriers had more pronounced age-related atrophy (Gonneaud et al. 2016; Bussy et al. 2019). Although not the primary interest of this paper, age-related *APOE4* effect was investigated in a subset of 203 genotyped participants (52 ADB-T1, 48 ADB-T2, 52 HA-T1 and 51 HA-T2) with models investigating the interaction of *APOE4* with age. No *APOE4* effect was found to be related to hippocampal volume with age. Therefore, *APOE4* was discarded in our following analyses.

#### 2.3.2 Impact of sequence type on the relationship between hippocampal subfield volumes with age

We studied the impact of the sequence type on the relationship between hippocampal subfield volumes with age (*3*). Here, we examined whether the intercept and/or slope of the predicted model was influenced by sequence-type after covarying by sex, ICV, and ipsilateral hippocampal GM or WM volume as fixed effects, and dataset and subjects as random effects.

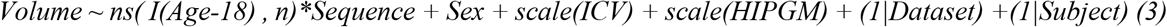

Figure 5 and Supplementary figure 11 illustrate the relationship between age and hippocampal subfield volumes when extracting from T2w and slab images, compared to when extracting using T1w data. To visualize that, we used model coefficients to predict T1, T2w, and slab subfield volumes divided by the predicted volume at age 18 for a subject of mean ICV, and mean ipsilateral hippocampal GM or WM volume extracted from T1w images. The goal of dividing by the predicted volume at age 18 extracted from T1w images, regardless of sequence, was to remove the potential effect of over- or under-estimation of a specific sequence and thus to focus on the age-related relationship differences. Supplementary figures 10 and 12 represent the significant over- or under-estimation of the volumes, in addition to the significant age-related relationships encountered when using solely model coefficients to predict T1, T2w, and slab subfield volumes.

**Figure 5:**
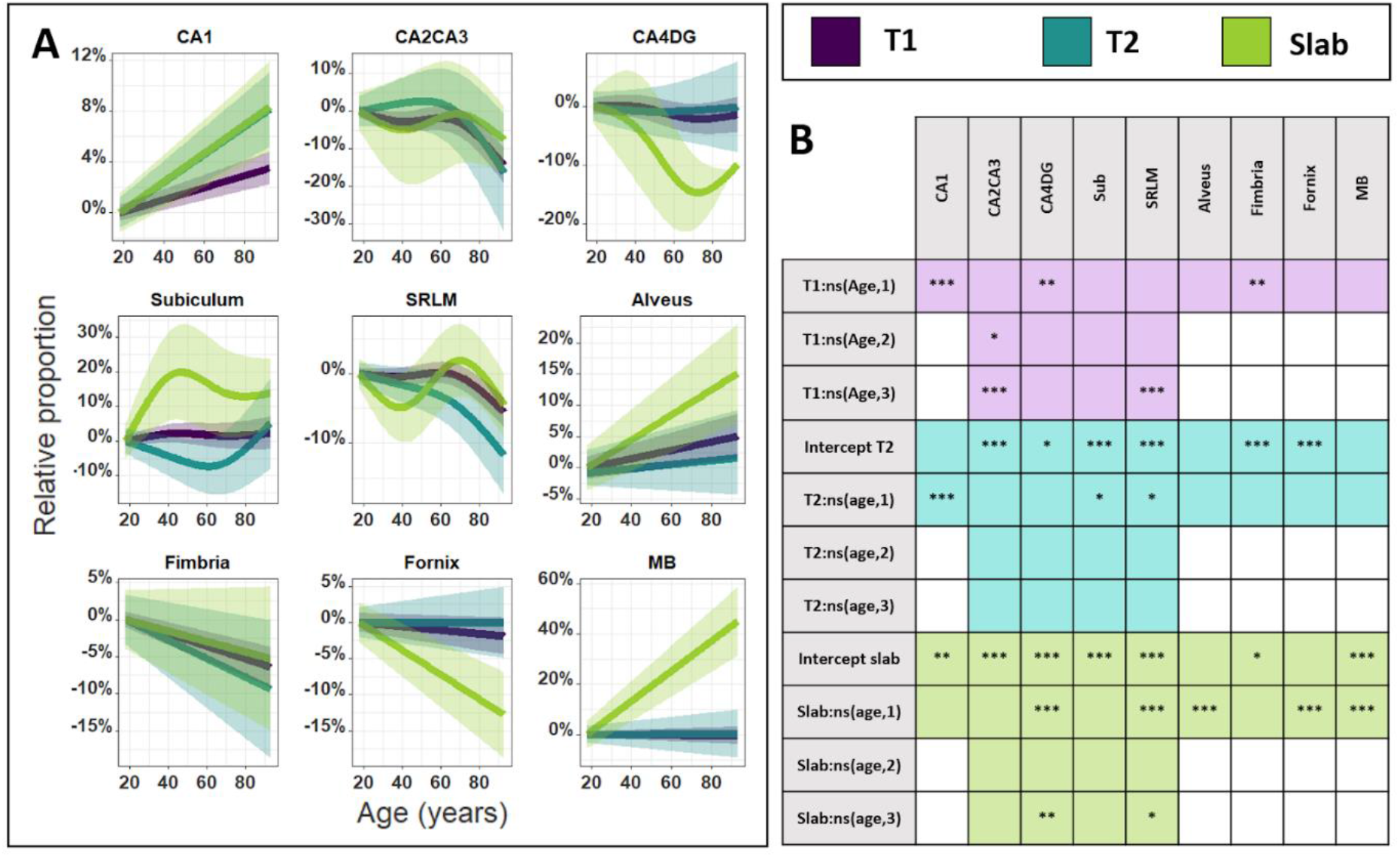
**A.** Estimated fixed effect plots showing the relationships, separated by sequence-type, between age and the relative proportion of the right hippocampal subfields, using the predicted volume at age 18 for a subject of mean ICV and mean ipsilateral hippocampal GM or WM volume extracted from T1w images as baseline. Best fit model displayed for each subfield covaried by ipsilateral hippocampal GM or WM volume, ICV, and sex as fixed effects and dataset, sequence, and subject as random effects. **B.** Table describing the significant coefficients for the different relationships with age and the intercepts. T1w was used as reference sequence in the model and demonstrated a significant linear increase for CA1 (p=2.65×10^−6^), decrease for fimbria (p=9.85×10^−3^) and third order decrease for CA2CA3 (p=6.59×10^−5^) and SRLM (p=1.33×10^−5^). Significant differences were found with slab compared to T1w, with the CA1 (p=0.0102) and MB (p=3.47×10^−14^), demonstrating a steeper increase with age, while CA4DG (p=1.93×10^−3^) and fornix (p=9.41×10^−7^) showing a steeper decline with age. Best fit models estimated from the T2w sequence exhibited similar relationships with age than the models obtained with T1w images except that T2w expressed steeper increases with age for CA1 (p=5.49×10^−4^). Supplementary figure 10 described in more details the significant intercept differences. Similar investigations made in the left hemisphere can be found in the Supplementary figures 11 and 12. * p<0.05; ** p<0.01 and *** p<0.001 after Bonferroni correction.

#### 2.3.3 Impact of sequence type on volume estimates

Given the use of various MRI parameters to study the hippocampus subfields in the literature, we used the test-retest dataset to compare the volume estimates from T1w, T2w, and slab sequences in the same participants to see how these diverse parameters impact subfield volumes estimation. A dependent 2-group Wilcoxon signed rank test was performed to compare the volume estimates from T1w, T2w, and slab sequences. Here, we used a Bonferroni correction significance level adjusted for 54 multiple comparisons (18 subregions x 3 sequence types), resulting in a significance level of p<0.00093. Intraclass correlation coefficients (ICCs;*psych_1.8.12* package) were determined to reflect the degree of consistency (ICC [3.1]) between the volume estimates of the different sequences. ICCs were interpreted according to previously established criteria: “excellent”: 1.00-0.75, “good”: 0.74-0.60, “fair”: 0.59-0.40, and “poor”: 0.39-0.00 (Cicchetti et al 1994). The mean percentage volume difference was determined to calculate the extent of the differences in the volume estimations for each sequence.

## Results

### 3.1. Relationship between hippocampal subfield volumes and age

#### 3.1.1. All datasets normalized by the ICV

After covarying for ICV and assessing linear, second- and third-order relationships with age using the AIC, second-order models demonstrated to be the best fit for bilateral GM and WM hippocampus (Figure 2): right HIPGM (p 3.33×10^4^), right HIPWM (p=1.05×10−^3^), left HIPGM (p2.27×10 ‘) and left HIPWM (p=1.18×10^−5^). These results show that, when accounting for head size difference, the relative volumes of the bilateral hippocampi were reduced by approximately 10% between age 18 and 93, with a steeper decline after age 50.

To test which subfields are involved in this 10% relative volume decrease of the hippocampi with age, individual subfield relationships with age were examined. Third-order models were observed to be the best fit for all hippocampal subfields bilaterally (Figure 3 and Supplementary figure 4). Significant monotonic decreases were found for the right CA2CA3 (p=3.55×10^−7^), CA4DG (p=4.28×10^−3^), SRLM (p=7.74×10^−7^), fimbria (p=9.45×10^−4^) and fornix (p=1.79×10^−4^). These results indicate that relative volumes of the CA2CA3, SRLM, fimbria and fornix decreased by approximately 20%, while CA4DG decreased by 10% between age 18 and 93. Interestingly, volumetric impairments seem to start earlier in CA4DG, SRLM and fimbria compared to the CA2CA3 and fornix, which seem relatively preserved until age 60. Similar relationships were found in the left hemisphere (Supplementary figures 6 and 7). In contrast, the bilateral CA1, subiculum, alveus and MB did not express significant relationships with age, and therefore were not implicated in the global volume decrease of the hippocampus.

#### 3.1.2. All datasets normalized by the hippocampal volume

In the right hemisphere, when covarying for ipsilateral hippocampal GM or WM volume and ICV, all hippocampal subfield volumes were shown to have a best fit third-order relationship with age. Significant monotonic increases were found for the relative proportion of the right CA1 (p=1.19×10^−10^). The relative volume of CA1 demonstrated a 7% increase between age 18 and 93. Significant monotonic decreases were found for the right CA2CA3 (p=3.19×10^−4^) and SRLM (p=5.29×10^−6^) (Figure 4 and Supplementary figure 5). These results indicate that relative volumes of the CA2CA3 decreased by approximately 15% and SRLM by 5% between age 18 and 93. No significant volume interaction with age was found for the right CA4DG, subiculum, alveus, fimbria, fornix and MB. Similar relationships were found in the left hemisphere (Supplementary figures 8 and 9).

#### 3.1.3. Impact of sequence on subfield volume relationship with age

T1w volumes demonstrated significant linear increases for CA1 (p=2.65×10^−6^), decreases for fimbria (p=9.85×10^−3^) and third-order relationships for CA2CA3 (p=6.59×10^−5^) and SRLM (p=1.33×10^−5^) highlighting a steeper decline after age 60 for these subfields (Figure 5A). Compared to T1w, significant differences were found with slab for the CA1 (p=0.0102) and MB (p=3.47×10^−14^), demonstrating a steeper increase with age, while CA4DG (p=1.93×10^−3^) and fornix (p=9.41×10^−7^) manifested a stronger decline with age. Volumes estimated from the T2w sequence exhibited similar relationships with age when compared to volumes estimated from T1w, except that T2w expressed steeper increase for CA1 (p=5.49×10^−4^). In Figure 5A, the different age relationships for each sequence are represented without intercept difference to emphasize the effect of age on the estimation. While not plotted, we see in Table 5B (and in more details in Supplementary figure 10) that slab intercepts were smaller for CA1, CA2CA3, subiculum, fimbria, and MB, while they were higher for CA4DG and SRLM. In addition, compared to T1w volumes, T2w volumes expressed smaller intercepts for CA2CA3, subiculum, and fimbria and higher intercepts for CA4DG, SRLM and fornix. Similar relationships were found in the left hemisphere (Supplementary figures 11 and 12).

### 3.2. Impact of different sequence on hippocampal subfield volumes estimates

We compared the volume estimates of the right hippocampal subfields obtained from T1w, T2w, and slab sequences within the same subjects (Figure 6). Significant differences were obtained between T1w and slab sequences for all the subfields except the SRLM. These results demonstrated that T1w images lead to larger volume estimates than the slab sequence, on average by 11.2% (Supplementary table 1), except for the CA4DG which is larger using slab images. T2w images appeared to render similar volumes compared to T1w images, except for the CA2CA3, which was larger when extracted from T1w. Moreover, T2w volumes estimates were on average 8.1% larger than the volumes extracted from slab sequences. These findings suggest that slab images may underestimate volumes related to other acquisitions.

**Figure 6:**
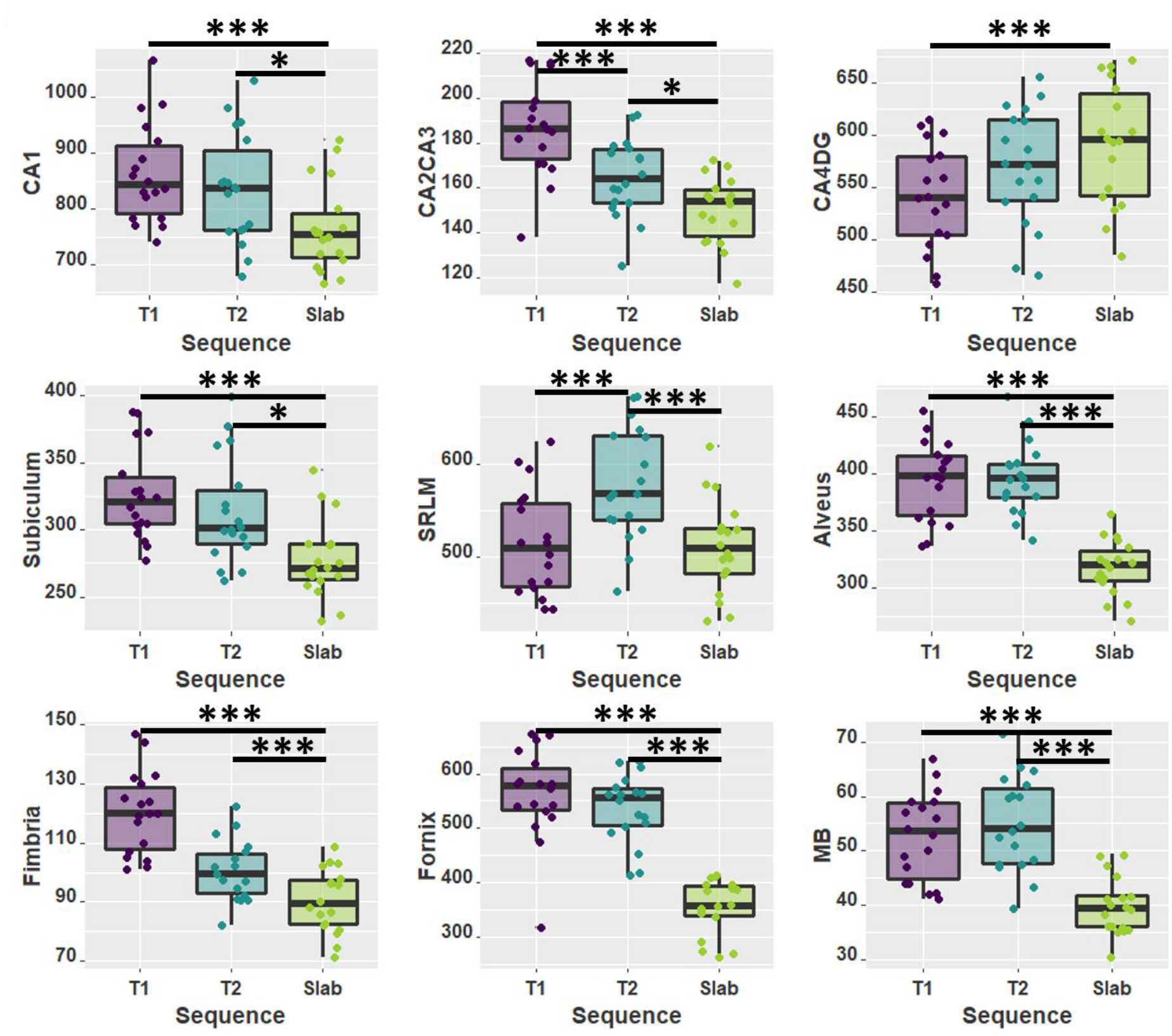
Boxplots illustrating the right volume estimates from T1w, T2w and slab sequences from the same participants as well as the dependent 2-group Wilcoxon signed rank test results (Supplementary table 1). Similar results were found in the left hemisphere (Supplementary figure 13). * p<0.05; ** p<0.01 and *** p<0.001 after Bonferroni correction for 54 comparisons (18 subfields x 3 sequence types).

Finally, ICC (3,1) was used to calculate the consistency between each dataset volume estimates (Figure 7). High ICC was mostly found between T1w and slab sequences. We can also observe that the left hippocampus revealed lower ICC in most subfields. Subregions exhibiting poor consistency included left fimbria and fornix.

**Figure 7:**
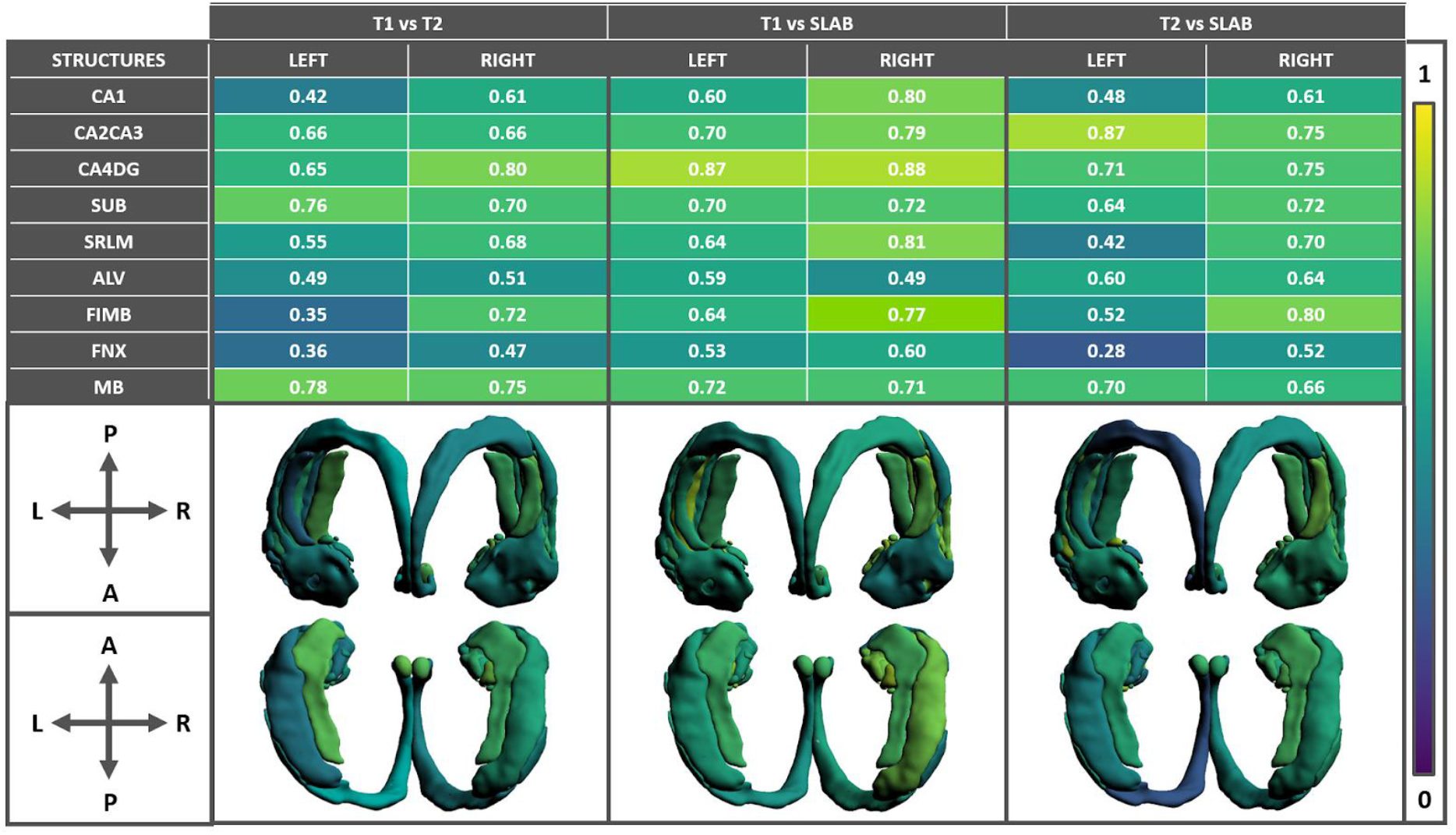
Representation of ICC consistency (3,1) of hippocampal subfields volume estimates from T1w, T2w, and slab images. Colour scale serves as an indicator of the ICC values: yellow for an ICC of 1 and dark blue for an ICC of 0.

## Discussion

The aim of this paper is to examine the hippocampal subfields and WM subregions throughout healthy aging. Normalized for ICV, we found that all subfields and WM subregions expressed a volumetric decrease with age with the exception of CA1, subiculum, alveus and MB. These findings are in contradiction with previous investigations that demonstrated a significant impact of age on the CA1 (Shing et al. 2011; Wisse et al. 2014; de Flores et al. 2015; Wolf et al. 2015; Daugherty et al. 2016). Although, other studies found no volumetric change with age for the CA1 (Voineskos et al. 2015) or even an increase with age for the CA1 and alveus when accounting for ipsilateral hippocampal volume (Amaral et al. 2018). No clear age-related relationship between the CA1-3 volumes with age was demonstrated in another study, with a significant decrease in the body but not in the head and tail (Malykhin et al. 2017). Regarding WM subregions, the alveus, fimbria and fornix have been previously mostly used as anatomical landmarks for the GM subfields definition (Mueller et al. 2007; La Joie et al. 2010; Wisse et al. 2012; Malykhin et al. 2017). In the present study, WM subregions were considered as an integral part of the hippocampal circuitry. CA1 and alveus were found to be stable with age which reproduces, using a larger sample, previous findings from our group (Amaral et al. 2018).

Researchers have consistently shown the CA1 subfield to be the first hippocampal subfield to be impacted in AD (Frisoni et al. 2008; de Flores et al. 2015; Adler et al. 2018). Therefore, we suggest that CA1 preservation in healthy aging could be of interest to identify early changes in hippocampus volume trajectories. SRLM has also been found to be especially impacted in patients with AD (Adler et al. 2018) as well as to a greater extent in *APOE4* carriers (Kerchner et al. 2014). Thus, it is interesting that our results also indicate a strong age-related atrophy in this subfield, although we did not find any *APOE4* effect. This could be in part because we did not have *APOE4* genotyping for a subset of our participants, preventing us from having enough data to fully investigate this genetic effect.

Complementary analyses were performed using ipsilateral hippocampus GM or WM volumes normalization. To our knowledge, this is the first study that analyzed the subfield volumetric relationship with respect to age while including global hippocampal atrophy. SRLM and fornix subregions expressed the highest relative volumetric impairment while the CA1 presented a relative volumetric preservation of its volumes with age. These findings replicate the results that we found using ICV normalization and provided a reasonable indication that the CA1 is a preserved hippocampal subfield in healthy aging. Dissimilarities with other publications could be explained by the variability of the hippocampal subfield atlases definition used in the literature (Yushkevich et al. 2015). We have previously found similar increases of the CA1 volumes (Amaral et al. 2018) after normalization using the same atlases and segmentation protocol in another cohort.

We hypothesize that the variability in findings within the literature may also be a function of the methodology used. For example, some groups approximated the hippocampal volumes using solely three contiguous slices (Daugherty et al. 2016) while others used nine slices (La Joie et al. 2010). Other studies have also expressed concern regarding the probable undersegmentation of CA1 and oversegmentation of the CA2-3 with FreeSurfer 5.3 (de Flores et al. 2015). Also, only a few studies previously examined the WM subregions of the hippocampus (Amaral et al. 2018; Malykhin et al. 2017), potentially because slab sequences do not always provide proper field-of-view to study the entire hippocampus circuitry.

Also, in the present study, we included participants aged from 18 to 93, an age range which is larger than what is used in most papers studying the effect of age in the hippocampus subfields (Mueller et al. 2007; Wolf et al. 2015; Dounavi et al. 2020). Almost all the significant relationships with age demonstrated third order relationship, often with an inflection point close to 60 years old. Similarly, a critical age in the acceleration of hippocampal degeneration was previously identified at 63 years old (Yang et al. 2013).

Mueller et al. (2018) compared different techniques to measure hippocampal subfield volumes and found that slab images were more sensitive to amyloid deposition and mild cognitive impairment status than whole brain T1w images. Also, slab images have demonstrated highly consistent results with those from T1w images and slightly better detection of group effects in atrophy rates between patients with cognitive impairment and controls (Das et al. 2012). In our study, we found that volumes extracted from slab images demonstrated different age-related relationships than the volumes extracted from T1w and T2w images. This could be explained by the relatively small number of participants having a slab image in our study. Indeed, 64% of our original slab scans had to be excluded due to high motion artifact or incomplete coverage of hippocampal GM and WM subregions (Mueller et al. 2018) compared to 43% for T1w and 25% for T2w (Supplementary figure 2). Of note, it is unclear how consistent quality control has been applied in other slab volumetric studies.

To our knowledge, our paper used, for the first time, a high-resolution isotropic whole brain T2w sequence with isotropic voxel dimensions of 0.64 mm. The main advantage is that this acquisition provides a voxel volume of 0.26 mm^3^ compared to 0.32 mm^3^ for the commonly used slab sequence or 1 mm^3^ for T1w sequence. Additionally, it allows for a clear delineation of the SRLM due to the hypo-intense contrast obtained in T2 contrast (Iglesias et al. 2015; Dounavi et al. 2020). This is of importance since it provides well-defined internal anatomical landmarks for segmentation protocols. T2w results primarily demonstrated similar results to T1w volume estimates but with the supplementary benefit of demonstrating higher precision of segmentation according to visual inspection of the labels.

The results presented in this paper should be interpreted with respect to several considerations and limitations. First, an important consideration is that this study aims to draw a global conclusion in how the different hippocampal subfields evolve across the lifespan using cross-sectional data. Thus, even though longitudinal analysis would be essential and more appropriate to assess the “true” relationship of the hippocampal subfield volumes and age, these studies require tremendous resources, time and dedication from the participants. Therefore, in the context of our research, since no longitudinal study used various sequences on healthy participants to study the hippocampus subfields, we used cross-sectional studies to approximate and study this topic.

Secondly, to create this study, we used datasets from multiple sites. This led to different inclusion criteria with regards to how each site defined healthy participants (Supplementary methods). Also, participants were scanned in different scanners, especially in the ADNI dataset, which itself included multiple sites. To counterbalance this issue, we performed rigorous preprocessing, quality control (Bedford et al. 2020), and statistical analyses to standardize the quality and the intensity of the images included in this study to the best of our ability.

In addition, the differing results with the slab scans must be interpreted carefully since this dataset was limited to 116 participants. Unfortunately, we did not have access to slab data from healthy participants across the entire lifespan. That is why we decided to include the test-retest dataset in order to increase the age-range of the slab sequence analyses. Thus, complementary analyses including more participants with a full coverage across the adult lifespan for the three modalities would be beneficial to validate these findings.

Another limitation relevant to the generalization of our findings is that our study is to a considerable extent dependent on our segmentation protocol (Chakravarty et al. 2013; Pipitone et al. 2014). Although MAGeT Brain has been validated by several studies (Pipitone et al. 2014; Makowski et al. 2018) and widely used to study the hippocampal subfields (Voineskos et al. 2015; Patel et al. 2017; Tardif et al. 2018; Patel et al. 2020), it remains that our subfield definitions are one among the multitude of atlases used in the literature. This highlights the importance of the work done by the Hippocampal Subfields Group (http://www.hippocampalsubfields.com/), which aims to standardize the definition of the medial temporal lobe segmentation, and to establish guidelines for subfield boundaries based on reference atlases and neuroanatomical landmarks visible postmortem and on MRI (Yushkevich et al. 2015; Olsen et al. 2019). A critical follow-up study would be to repeat this study to compare the effect of the MRI sequence using the complete and validated segmentation protocol described by the Hippocampal Subfield Group.

In light of our findings, we propose some guidelines to consider for future hippocampal subfields studies. For researchers interested in subtle relationships with age, we advocate the use of large age-range datasets since we demonstrated that most of our age-related relationships were non-linear with an inflection point occurring at approximately age 60. Slab anisotropic scans seemed to find different age-effects compared to those found with T1w and T2w isotropic images. Also, because of the characteristics of slab scans, we would suggest to be especially careful in the volume approximation of small structures, such as CA2CA3, fornix or MB, since these structures can be locally thinner than 2 mm. Moreover, we advise researchers using slab datasets to keep in mind that it is likely that in general the subfield volumes are underestimated compared to the estimation from standard T1w images (Das et al. 2012). Finally, for new scanning protocols, we encourage researchers to consider using high-resolution T2w sequences, which have shown promising sensitivity and accuracy. T2w images also demonstrated smaller volumes in most of the subfields compared to T1w images, but we predict that they may better estimate the “true” volumes since they have higher isotropic resolution and added contrast in key areas. T2w images rendered the same age-related relationships as T1w in most structures, although stronger age-related relationships were identified for the CA1 when compared to T1w images. Furthermore, high-resolution T2w images have demonstrated high-quality segmentation from visual inspection compared to slab and T1w scans (Figure 1), and provide whole brain images that permit hippocampal WM subregion investigation.

To conclude, we compared a wide range of datasets attempting to elucidate the relationship between hippocampal subfields with age. Certain subfields appeared to show reliable and reproducible relationships with age across various datasets. Nonetheless, the sensitivity of slab images to age-related changes in the hippocampus subfields deserves further exploratory analysis.

## Supplementary material

### 1) Participants

#### HA dataset

Here, we recruited 112 healthy individuals aged 18 to 80 and composed of 53 males and 58 females. The study was approved by the Research Ethics Board of the Douglas Mental Health University Institute in Montreal, Canada and written informed consent from all participants was obtained. Individuals with a history of PTSD, ADD/ADHD, bipolar disorder, schizophrenia, Alzheimer’s, Parkinson’s disease, physical injuries such as head trauma and concussion, alcohol or substance abuse were excluded. Participants with a Mini Mental State Examination (MMSE) score between 24-30 were considered eligible.

#### ADB dataset

ADB study evaluates volumetric differences in the architecture of brain circuitry in healthy seniors, seniors with mild cognitive impairment (MCI), and seniors with Alzheimer’s disease (AD). For the purpose of the present study, we only included 66 healthy seniors, aged 56-81, 27 males and 41 females. We used the same inclusion and exclusion criteria than for the HA cohort for the recruitment of this sample.

#### ADNI dataset

ADNI is a publicly available longitudinal multicenter study which began in 2004 and was created to examine the progression of AD. Here we focused on the ADNI-3 cohort (Weiner et al. 2017) and included cross-sectional information from 317 healthy participants, aged 56-95, 118 males and 199 females. To be included in the ADNI cohort, participants must have MMSE scores between 24-30, a CDR of 0 and be non-depressed, non-MCI, and nondemented. Participants with any significant neurologic disease, evidence of infection, infarction, or other focal lesions or with a history of alcohol or substance abuse were excluded.

#### Cam-CAN dataset

Cam-CAN is a large-scale cross-sectional research project from the University of Cambridge, England and was launched in October 2010. Epidemiological, cognitive, and neuroimaging data were acquired to understand how individuals can best retain cognitive abilities into old age. 652 healthy individuals, aged 18-88, 322 males and 330 females were included in our study. Participants were required to be cognitively healthy (MMSE 24-30), to meet hearing, vision, and English language ability criteria necessary for completing experimental tasks, and to be free of MRI or MEG contraindications and neurological or serious psychiatric conditions.

#### Test-retest dataset

The purpose of the test-retest dataset is to compare the volume estimates of the hippocampal subfields using different sequences of acquisition within the same subjects. Eighteen participants free of neurological disorders, aged 20-42 with a mean age of 26.9 including 7 males and 11 females were recruited in Montreal, Quebec, Canada.

**Supplementary figure 1:**
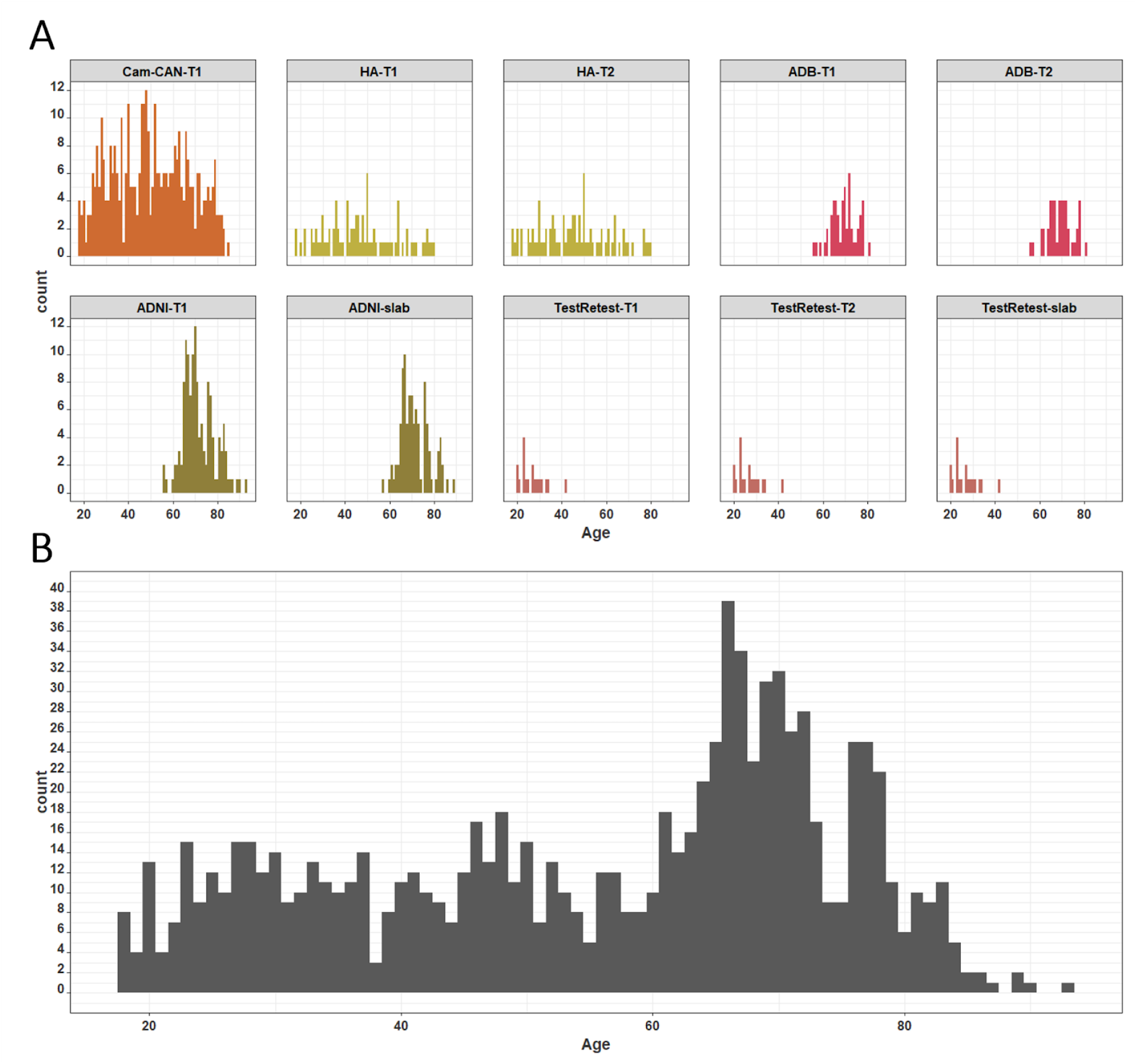
**A)** Age distribution by dataset and sequence type, **B)** Age distribution of the entire dataset.

### 2) Acquisition parameters

- T1w 3D MPRAGE : TR=2300 ms, TE=2.01 ms, TI=900 ms, resolution=1 mm isotropic, FOV=192 x 240x 192 mm, flip angle=9 degrees, GRAPPA factor = 2, echo spacing=7.4 ms, bandwidth=240 Hz/Px.
- High Resolution T2w 3D SPACE: TR=2500 ms, TE=198 ms, resolution = 0.64 mm, FOV =263 x 350 x 350 mm, CAIPIRINHA imaging, GRAPPA acceleration factor=2, bandwidth=625 Hz/Px, echo train duration=483 ms, turbo factor=143.
- Slab T2w 2D TSE: TR=8020 ms, TE=76 ms, slice thickness=2 mm, FOV=150 x 150 x 60 mm, flip angle=150 degrees, base resolution=384, 30 slices, interleaved slice acquisition, echo spacing=15.3 ms, bandwidth=107 Hz/Px, echo train per slice=48, turbo factor=8.

**Supplementary figure 2:**
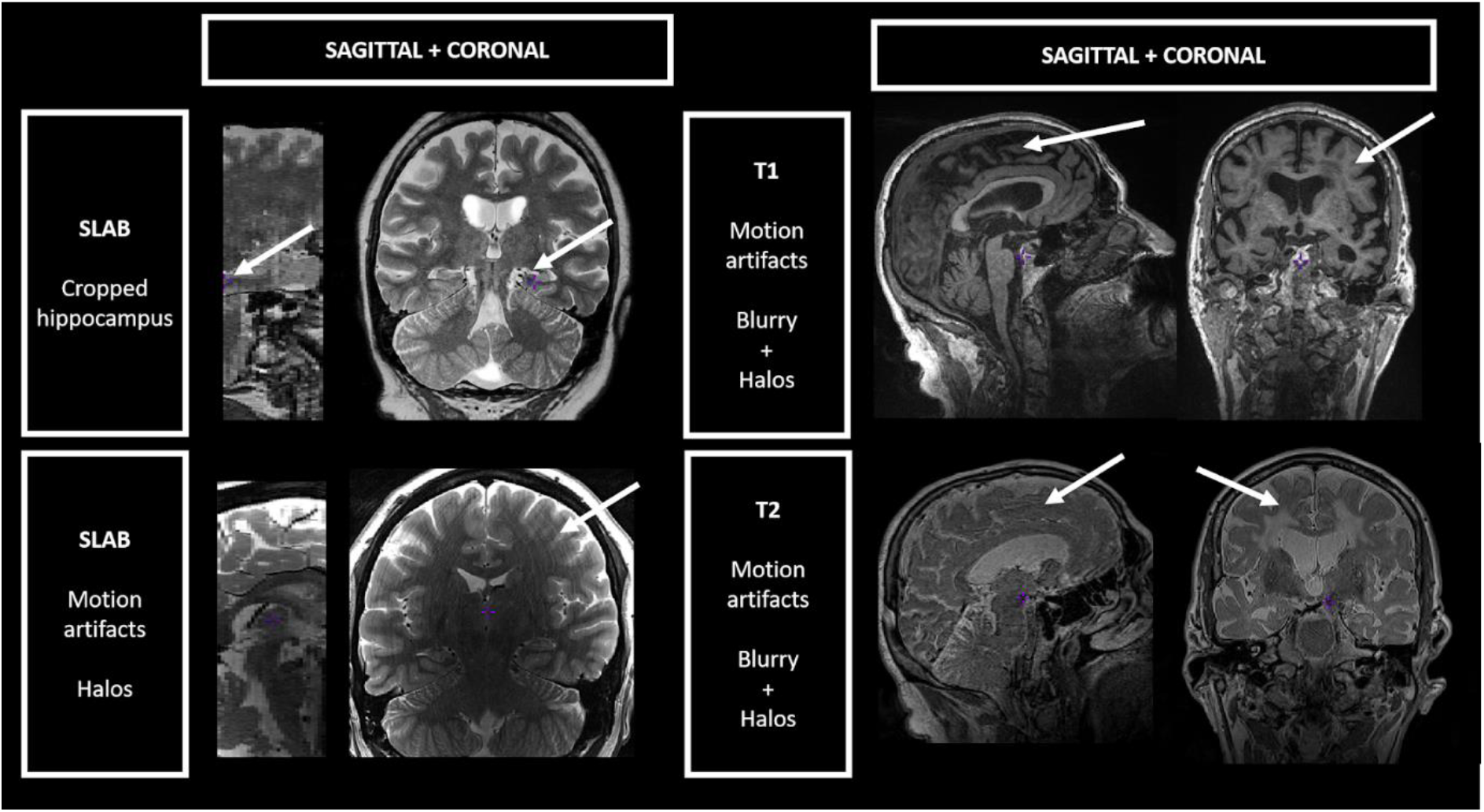
Examples of scans discarded due to inappropriate field-of-view or high motion artifacts. White arrows were added to indicate scan problems.

## Supplementary results

**Supplementary figure 3:**
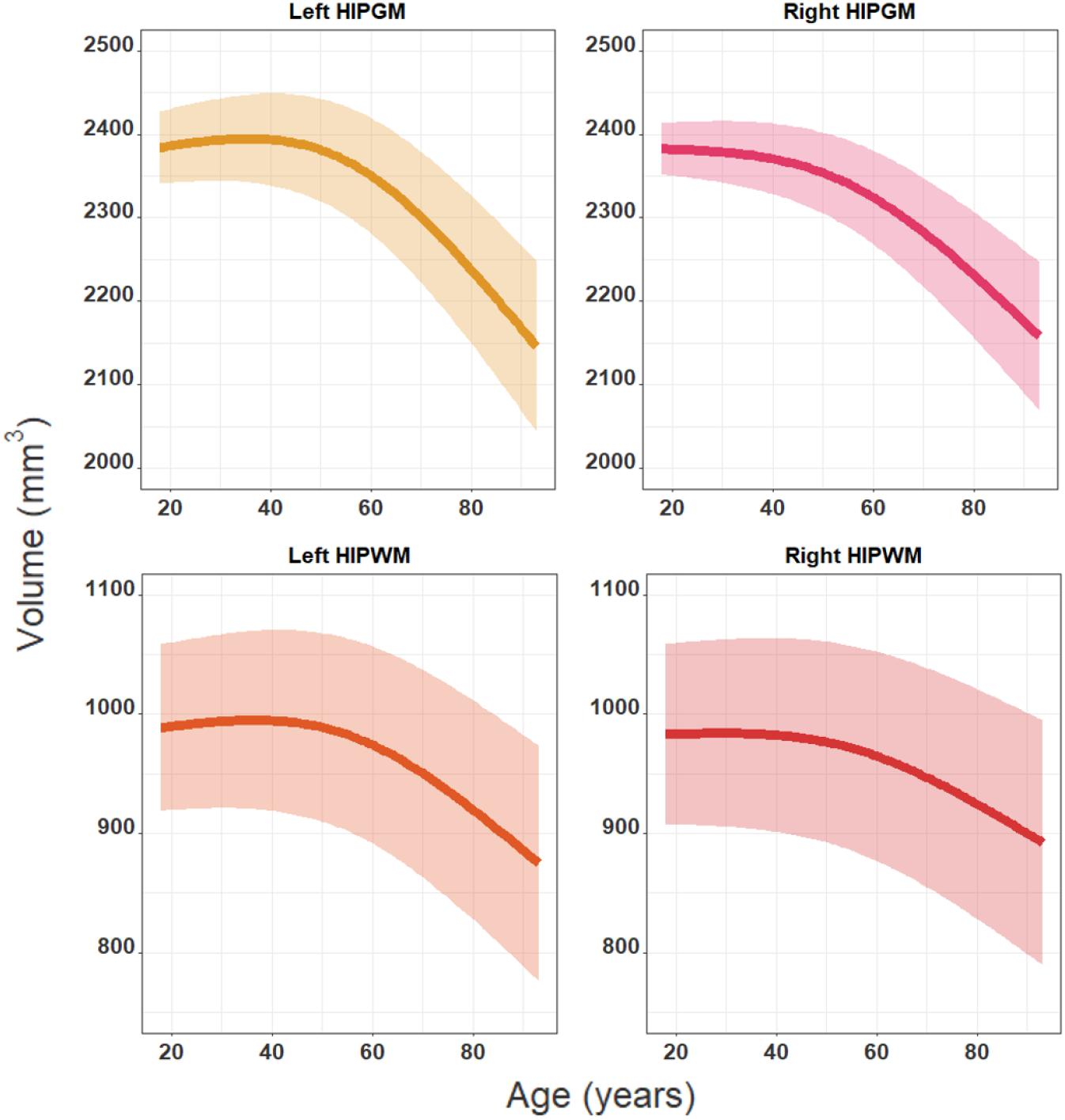
Best fit models showing the relationships between age and the hippocampal volume covaried by ICV and sex as fixed effects and dataset, sequence, and subjects as random effects. Second order relationships were found to be the best fit model for all the structures: right HIPGM (p=3.33×10^−4^), right HIPWM (p=1.05×10^−3^), left HIPGM (p 2.27×10^−5^) and left HIPWM (p 1.18×10^−5^).

**Supplementary figure 4:**
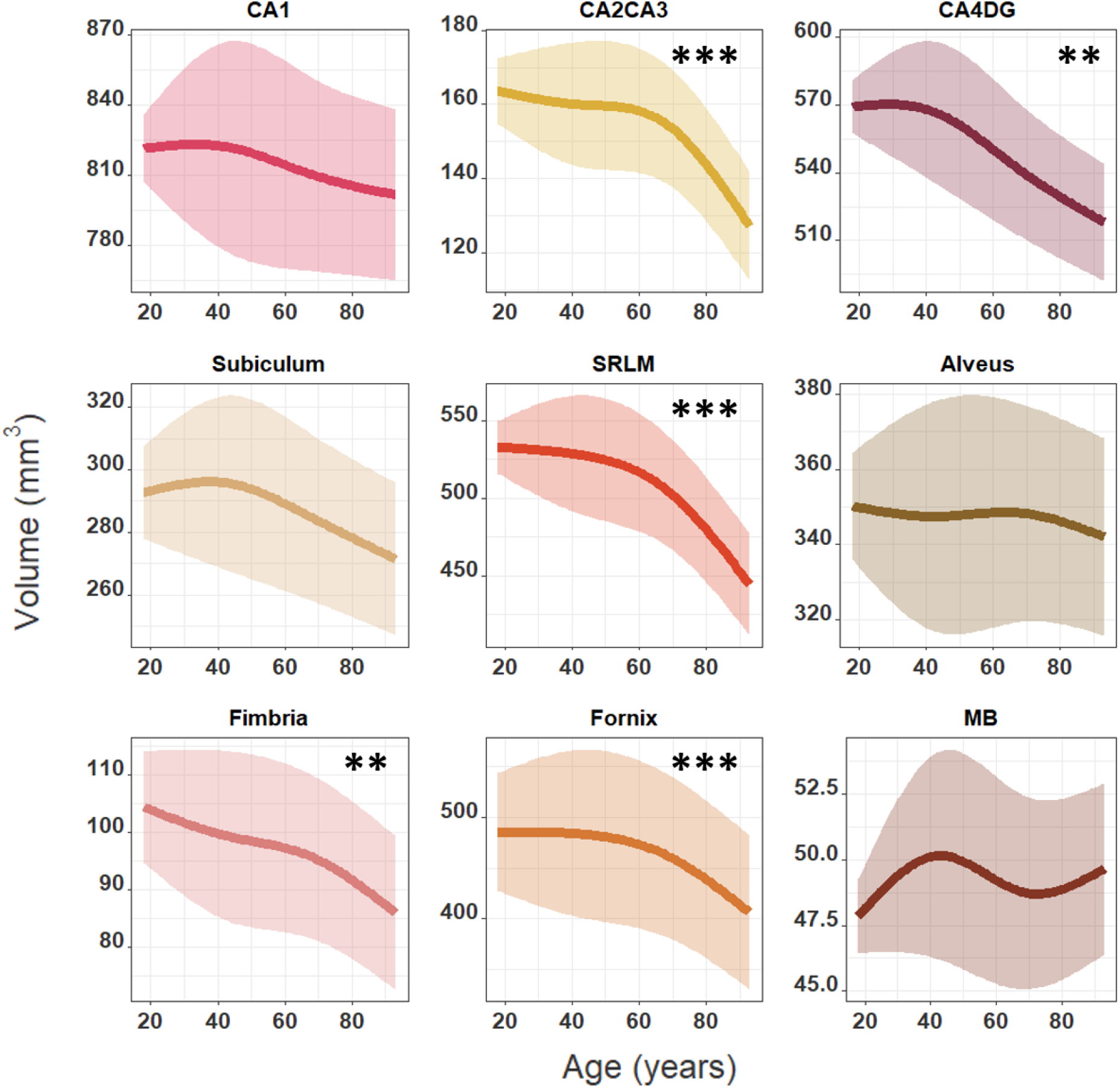
Best fit models showing the relationships between age and the volume of the right hippocampal subfields. Best fit models displayed for each subfield covaried by ICV and sex as fixed effects and dataset, sequence, and subject as random effects. Significant monotonic decreases were found for the right CA2CA3 (p=8.42×10^−7^), CA4DG (p=0.0085), SRLM (p=1.60×10^−6^), fimbria (p=0.0024) and fornix (p=4.79×10^−4^). Similar relationships were found in the left hemisphere (Supplementary figure 7). * p<0.05; ** p<0.01 and *** p<0.001 after Bonferroni correction.

**Supplementary figure 5:**
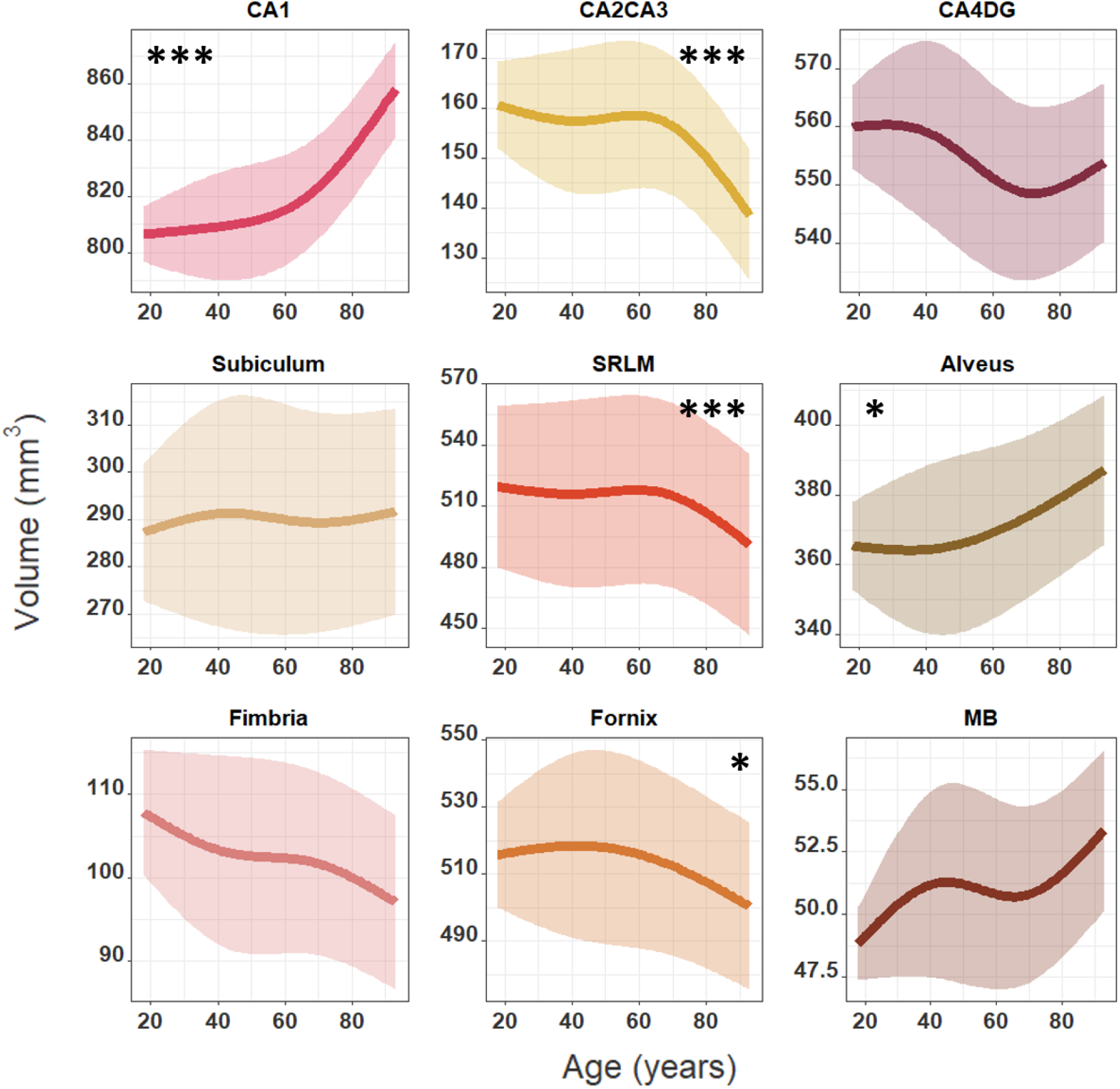
Best fit models showing the relationships between age and the volume of the right hippocampal subfields. Best fit models displayed for each subfield covaried by right hippocampal GM or WM volume, ICV and sex as fixed effects and dataset, sequence, and subject as random effects. Significant monotonic increases were found for the right CA1 (p=1.37×10^−10^) and alveus (p=0.010) and significant monotonic decreases were found for the right CA2CA3 (p=5.11×10^−4^), SRLM (p=7.99×10^−7^) and fornix (p=0.034). Similar relationships were found in the left hemisphere (Supplementary figure 9). * p<0.05; ** p<0.01 and *** p<0.001 after Bonferroni correction.

**Supplementary figure 6:**
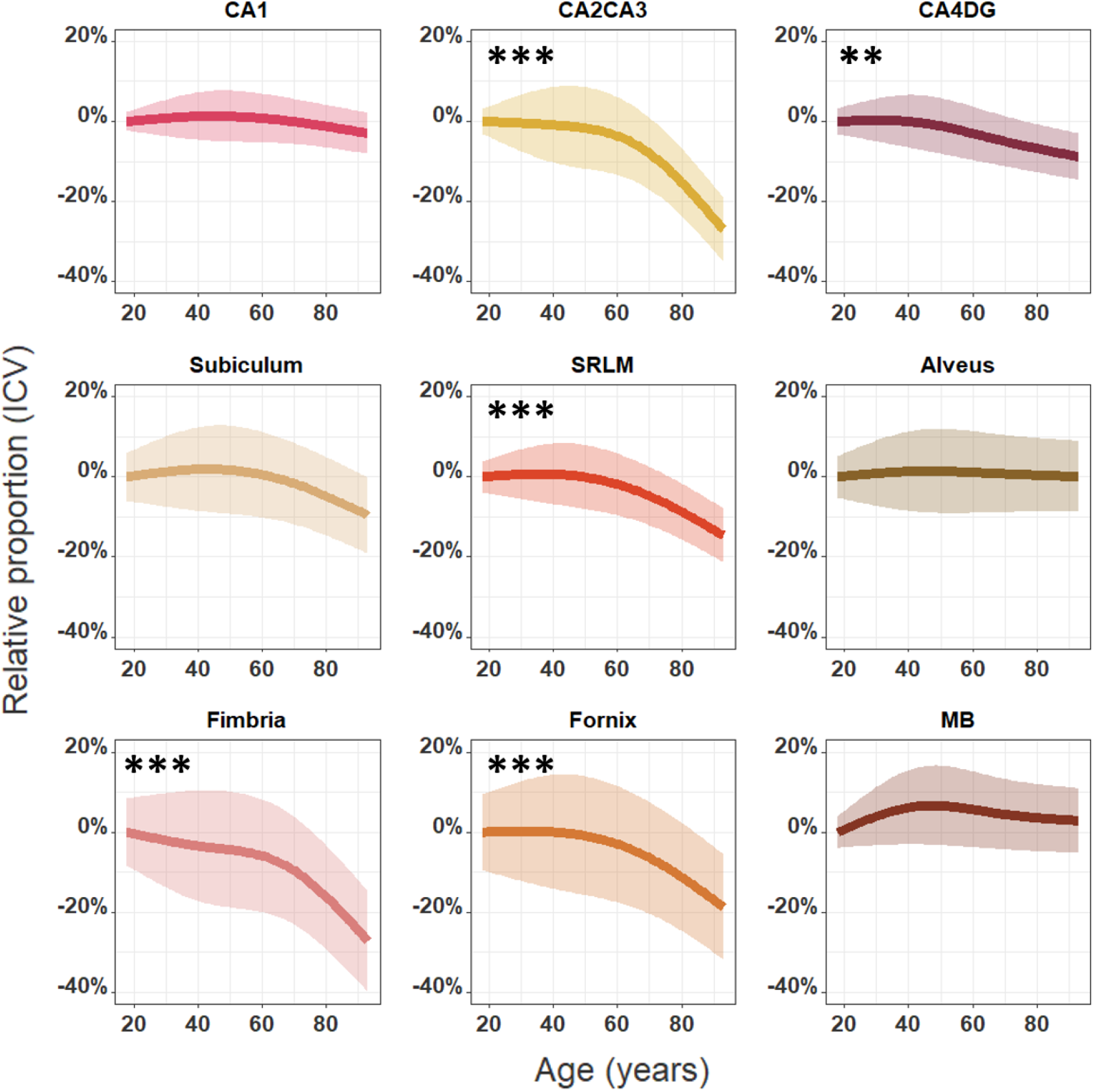
Best fit models showing the relationships between age and the relative proportion of the left hippocampal subfields, using the predicted volume at age 18 for a subject of mean ICV as baseline (same model with volume instead of relative proportion in Supplementary figure 7). Best fit models displayed for each subfield covaried by ICV and sex as fixed effects and dataset, sequence, and subject as random effects. Significant monotonic decreases were found for the left CA2CA3 (p=4.5×10^−6^), CA4DG (p=0.012), SRLM (p=3.60×10^−6^), Fimbria (p=6.16×10^−7^) and fornix (p=2.20×10^−5^). * p<0.05; ** p<0.01 and *** p<0.001 after Bonferroni correction.

**Supplementary figure 7:**
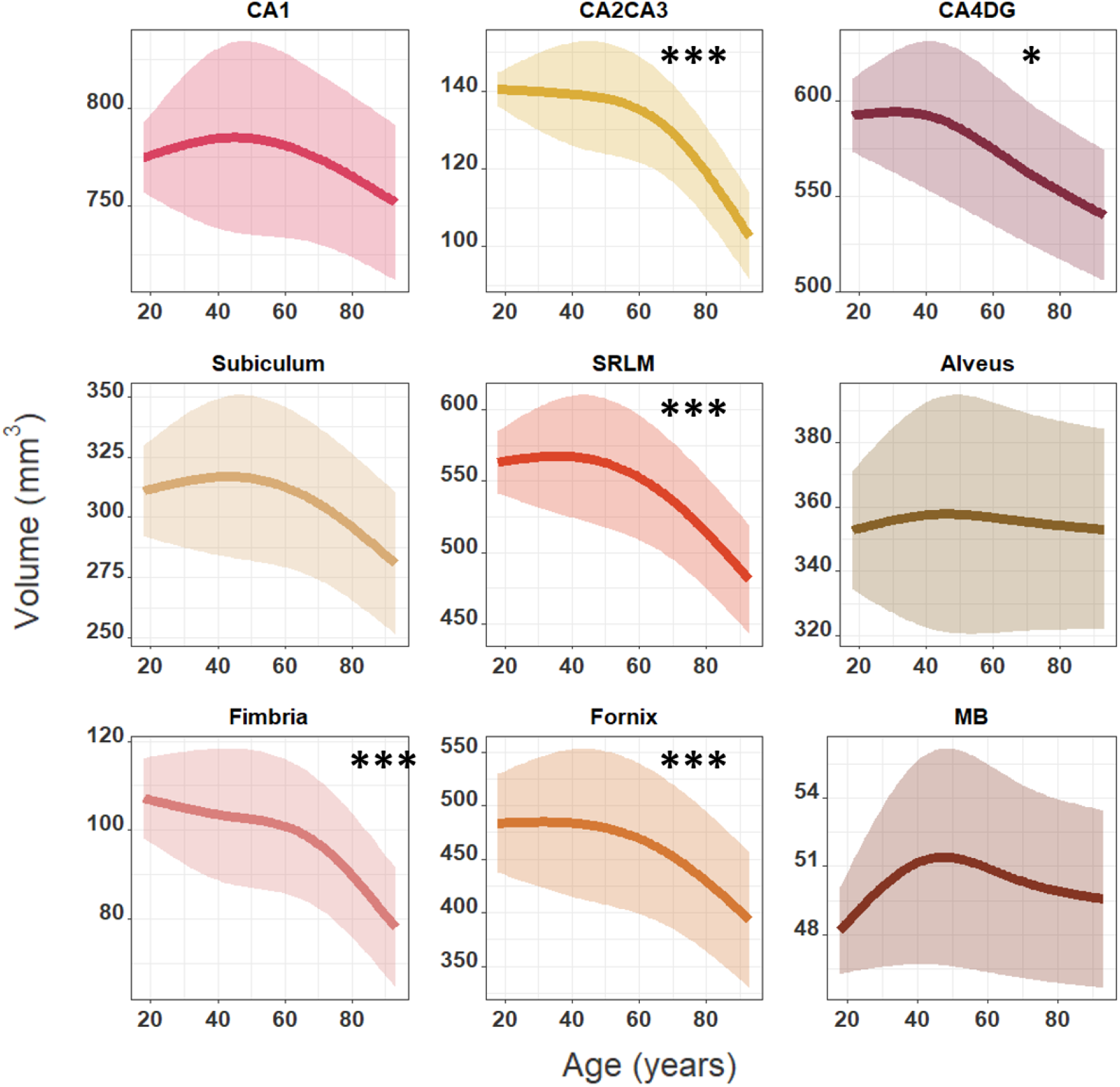
Best fit models showing the relationships between age and the volume of the left hippocampal subfields. Best fit models displayed for each subfield covaried by ICV and sex as fixed effects and dataset, sequence, and subject as random effects. Significant monotonic decreases were found for the left CA2CA3 (p=4.5×10^−6^), CA4DG (p=0.012), SRLM (p=3.60×10^−6^), fimbria (p=6.16×10^−7^) and fornix (p=2.20×10^−5^). * p<0.05; ** p<0.01 and *** p<0.001 after Bonferroni correction.

**Supplementary figure 8:**
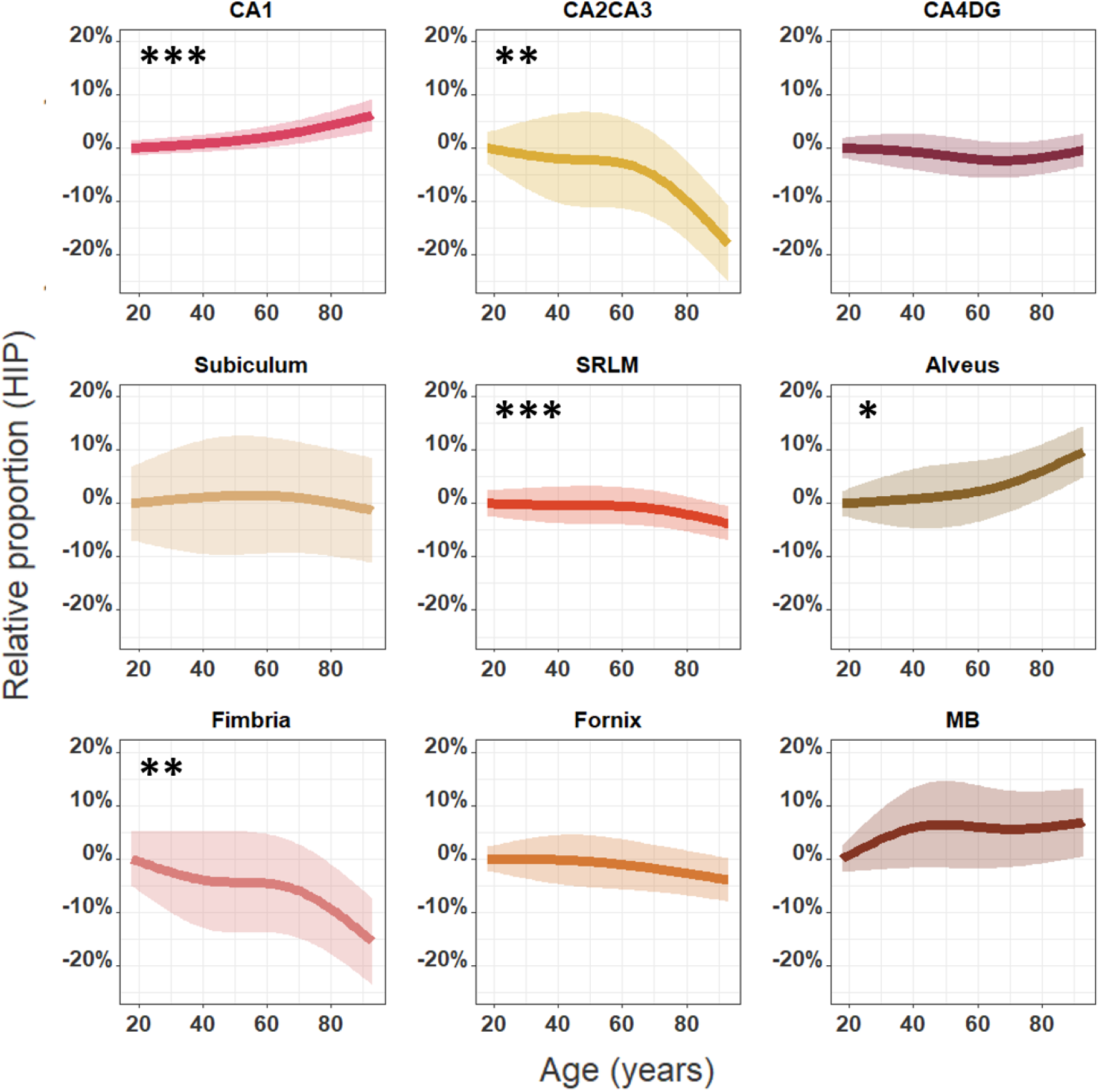
Best fit models showing the relationships between age and the relative proportion of the left hippocampal subfields, using the predicted volumes at age 18 for a subject of mean ICV and mean left hippocampal GM or WM volume as baseline (same model with volume instead of relative proportion in Supplementary figure 9). Best fit models displayed for each subfield covaried by left hippocampal GM or WM volume, ICV and sex as fixed effects and dataset, sequence, and subject as random effects. Significant monotonic increases were found for the right CA1 (p=1.62×10^−9^) and alveus (p=0.0135) and significant monotonic decreases were found for the right CA2CA3 (p=3.52×10^−3^), SRLM (p=7.56×10^−4^) and fimbria (p=0.0031). * p<0.05; ** p<0.01 and *** p<0.001 after Bonferroni correction.

**Supplementary figure 9:**
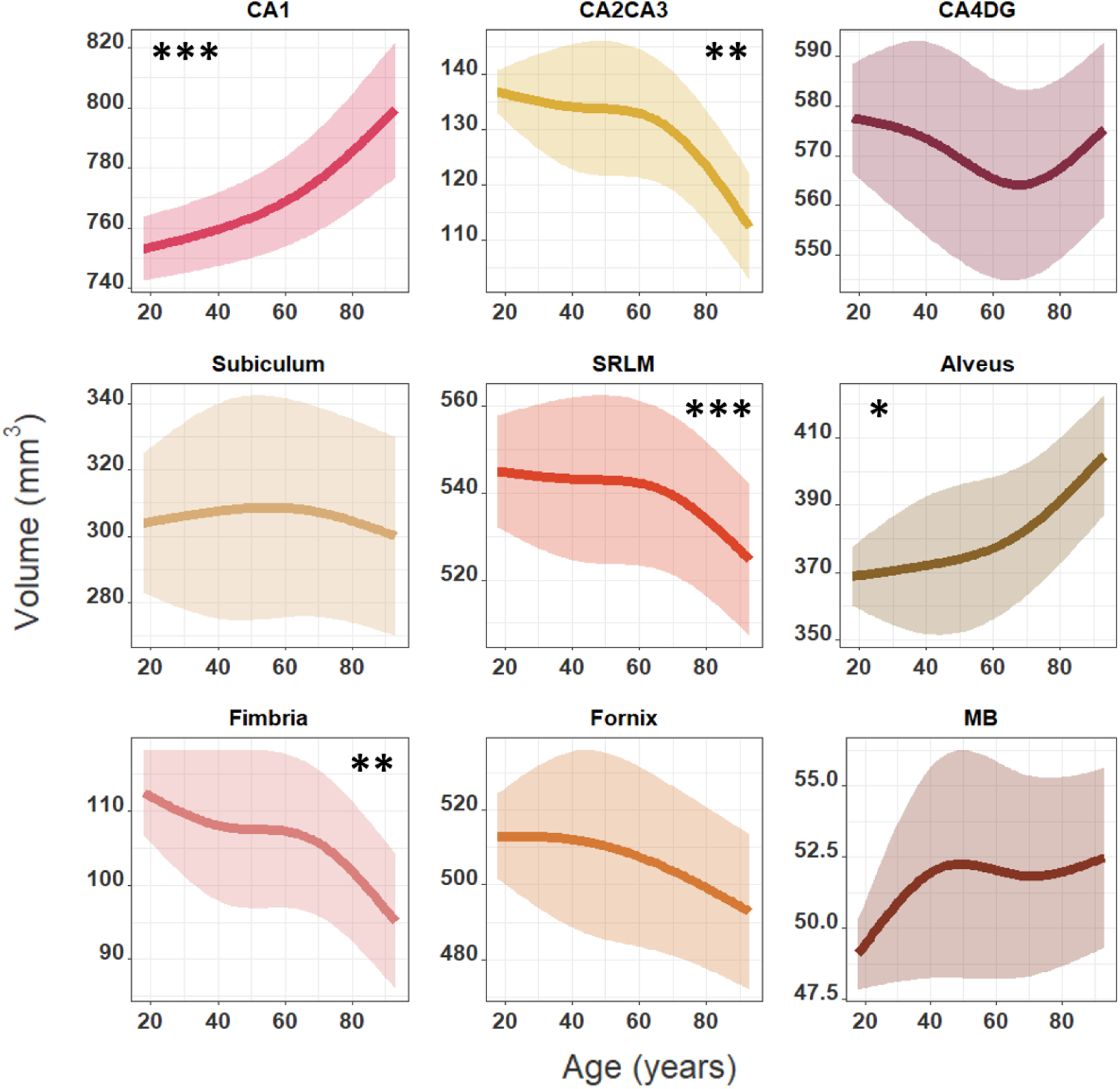
Best fit models showing the relationships between age and the volume of the left hippocampal subfields. Best fit models displayed for each subfield covaried by left hippocampal GM or WM volume, ICV and sex as fixed effects and dataset, sequence, and subject as random effects. Significant monotonic increases were found for the right CA1 (p=1.62×10^−9^) and alveus (p=0.0135) and significant monotonic decreases were found for the right CA2CA3 (p=3.52×10^−3^), SRLM (p=7.56×10^−4^) and fimbria (p=0.0031). * p<0.05; ** p<0.01 and *** p<0.001 after Bonferroni correction.

**Supplementary figure 10:**
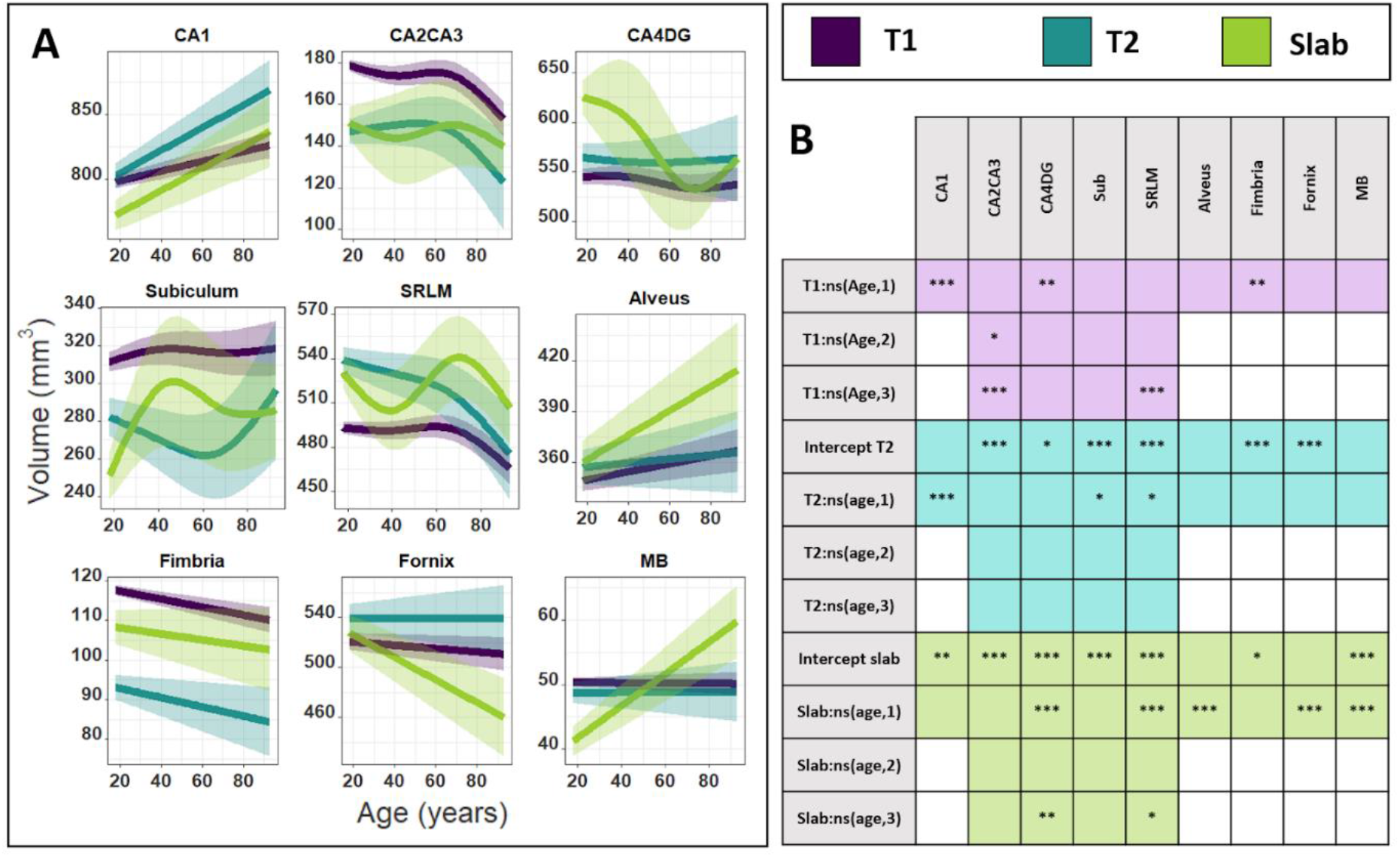
**A.** Estimated fixed effect plots showing the relationships, separated by sequence-type, between age and right hippocampal subfield volumes.. Best fit models displayed for each subfield covaried by right hippocampal GM or WM volume, ICV and sex as fixed effects and dataset, sequence, and subject as random effects. **B.** Table describing the significant coefficients for the different relationships with age and the intercepts. T1w was used as reference sequence in the model and demonstrated a significant linear increase for CA1 (p=2.65×10^−6^), decrease for fimbria (p=9.85×10^−3^) and third order decrease for CA2CA3 (p=6.59×10^−5^) and SRLM (p=1.33×10^−5^). Significant differences were found with slab compared to T1w, with the CA1 (p=0.0102) and MB (p=3.47×10^−14^), demonstrating a steeper increase with age, while CA4DG (p=1.93×10^−3^) and fornix (p=9.41×10^−7^) showing a steeper decline with age. Best fit models estimated from the T2w sequence exhibited similar relationships with age than the models obtained with T1w images, except that T2w expressed steeper increase with age for CA1 (p=5.49×10^−4^). Furthermore, slab demonstrated significant smaller intercepts for CA1, CA2CA3, subiculum, fimbria, MB and larger intercepts for CA4DG and SRLM. T2w demonstrated significant smaller intercepts for CA2CA3, subiculum, fimbria and larger intercepts for CA4DG, SRLM and fornix Similar investigations made in the left hemisphere can be found in Supplementary figures 11 and 12. * p<0.05; ** p<0.01 and *** p<0.001 after Bonferroni correction.

**Supplementary figure 11:**
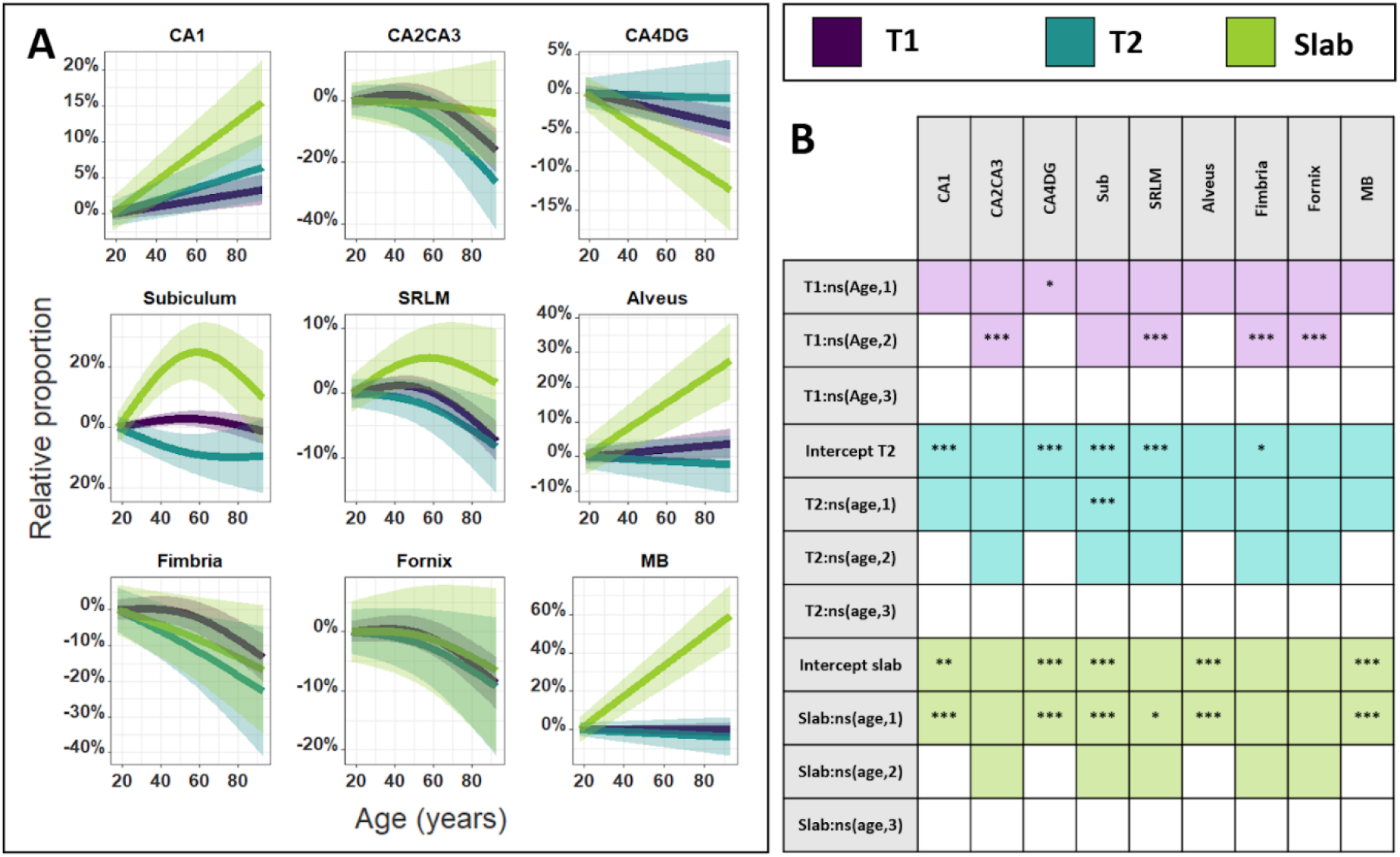
**A.** Estimated fixed effect plots showing the relationships, separated by sequence-type, between age and the relative proportion of the left hippocampal subfields, using the predicted volume at age 18 for a subject of mean ICV and mean left hippocampal GM or WM volume extracted from T1w images as baseline. Best fit model displayed for each subfield covaried by left hippocampal GM or WM volume, ICV, and sex as fixed effects and dataset, sequence, and subject as random effects. **B.** Table describing the significant coefficients for the different relationships with age and the intercepts. T1w was used as reference sequence in the model and demonstrated a significant linear decrease for CA4DG (0.0258), a second order decrease for CA2CA3 (p=7.71×10^−4^), SRLM (p=3.00×10^−5^), fimbria (p=2.99×10^−4^) and fornix (p=7.29×10^−4^). Significant differences were found with slab compared to T1w, with the CA1 (p=1.84×10^−6^), alveus (p=1.11×10^−9^) and MB (p=3.6×10^−16^) demonstrating a steeper increase with age, while CA4DG (p=1.91×10^−4^) showing a steeper decline with age. Best fit models estimated from the T2w sequence exhibited similar relationships with age than the models obtained with T1w images. Supplementary figure 12 described in more details the significant intercept differences. * p<0.05; ** p<0.01 and *** p<0.001 after Bonferroni correction.

**Supplementary figure 12:**
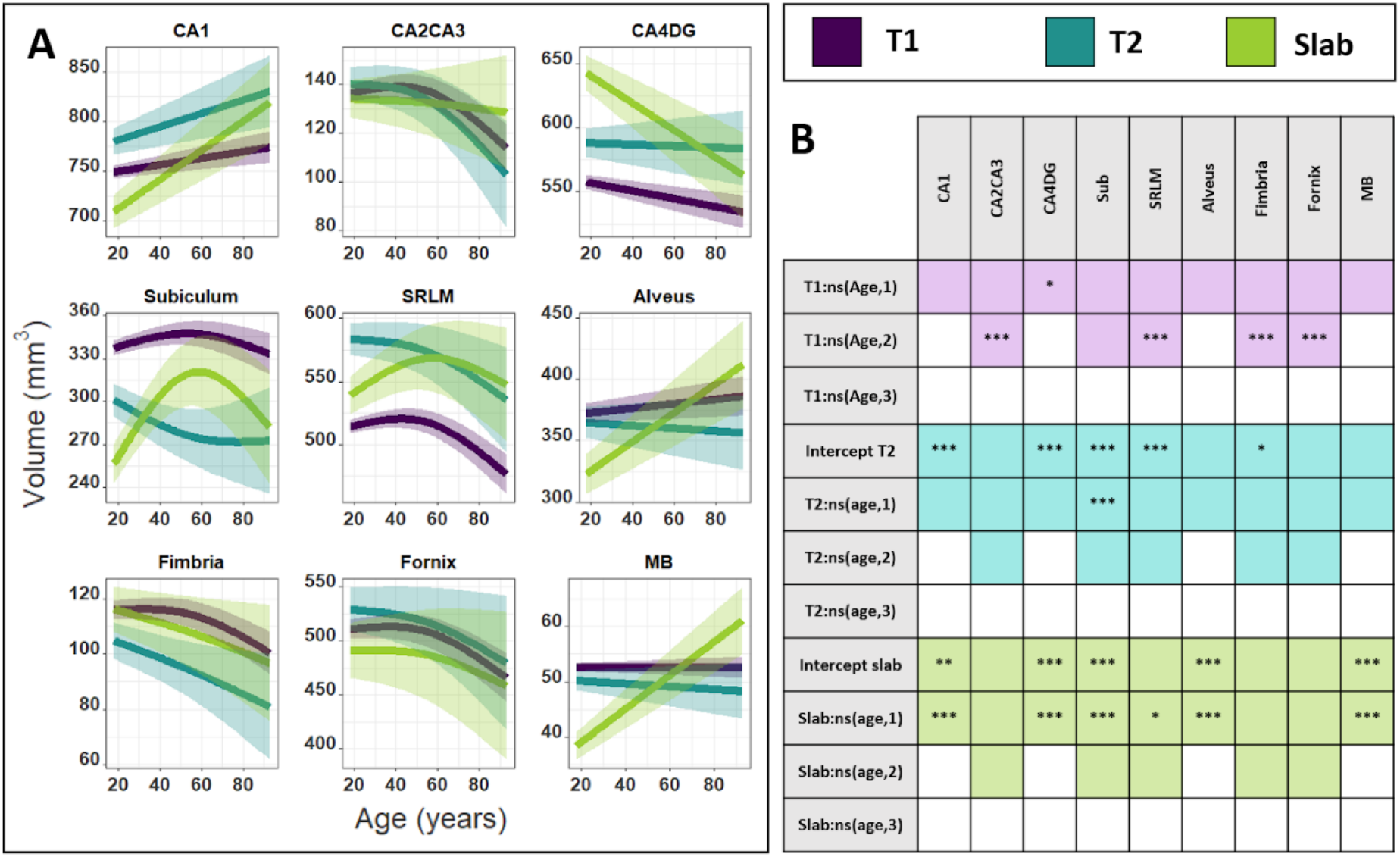
**A.** Estimated fixed effect plots showing the relationships, separated by sequence-type, between age and right hippocampal subfield volumes.. Best fit models displayed for each subfield covaried by ipsilateral hippocampal GM or WM volume, ICV and sex as fixed effects and dataset, sequence, and subject as random effects. **B.** Table describing the significant coefficients for the different relationships with age and the intercepts. T1w was used as reference sequence in the model and demonstrated a significant linear decrease for CA4DG (0.0258), a second order decrease for CA2CA3 (p=7.71×10^−4^), SRLM (p=3.00×10^−5^), fimbria (p=2.99×10^−4^) and fornix (p=7.29×10^−4^). Significant differences were found with slab compared to T1w, with the CA1 (p=1.84×10^−6^), alveus (p=1.11×10^−9^) and MB (p=3.6×10^−16^) demonstrating a steeper increase with age, while CA4DG (p=1.91×10^−4^) showing a steeper decline with age. Furthermore, slab demonstrated significant smaller intercepts for CA1, subiculum, alveus, MB and larger intercepts for CA4DG. T2w demonstrated significant smaller intercepts for subiculum and fimbria and larger intercepts for CA4DG and SRLM. * p<0.05; ** p<0.01 and *** p<0.001 after Bonferroni correction.

**Supplementary figure 13:**
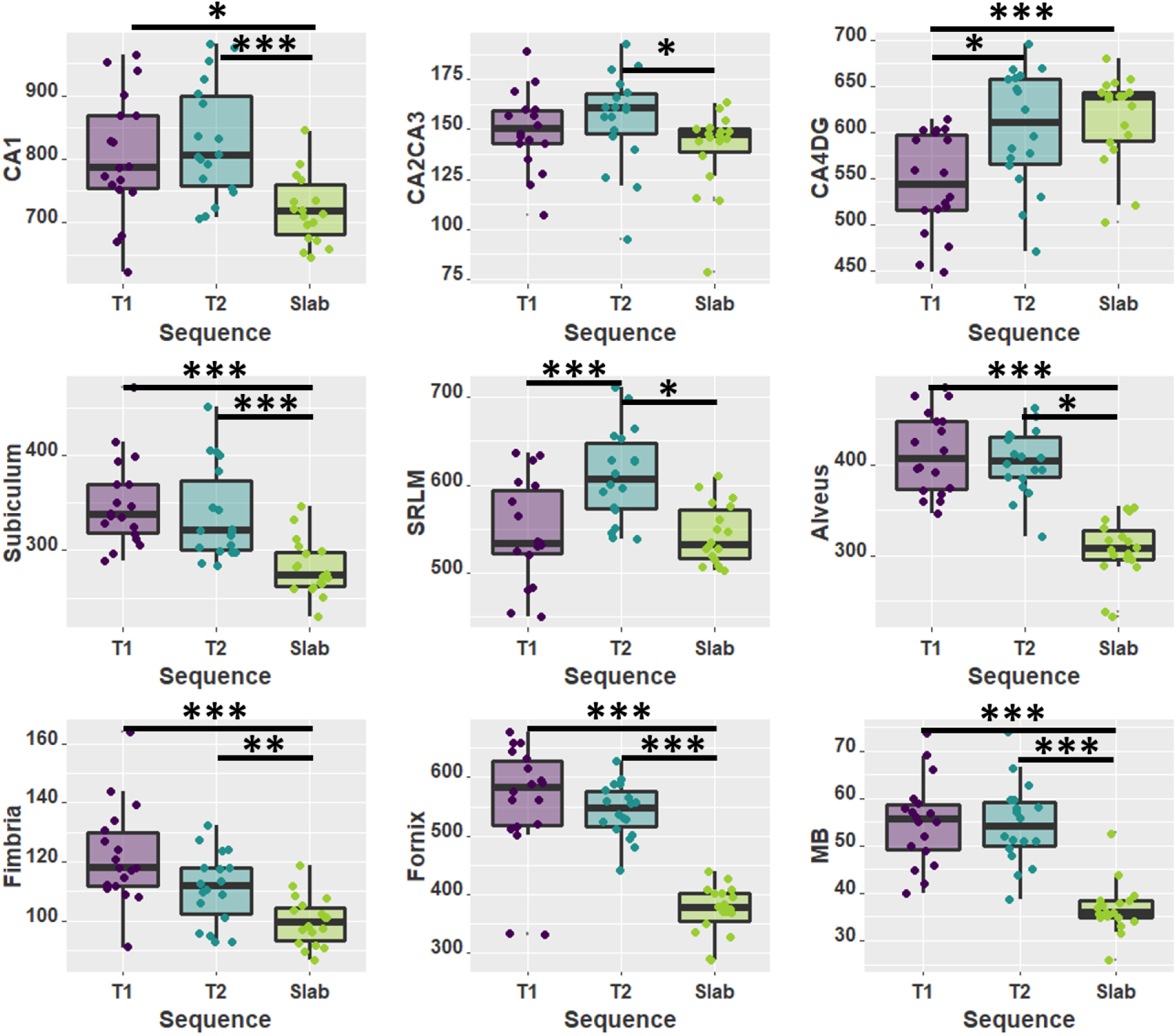
Boxplot comparing the left volume estimates from T1w, T2w and slab sequences from the same participants. Statistical tests performed using dependent 2-group Wilcoxon signed rank test. * p<0.05; ** p<0.01 and *** p<0.001 after Bonferroni correction for 54 comparisons (18 subfields x 3 sequence types).

**Supplementary table 1:** Percentage of volume difference between each sequence type and p-values obtained with dependent 2-group Wilcoxon signed rank test used to compare the volume estimates from T1w, T2w and slab sequences. Bonferroni correction adjusted for 54 multiple comparisons (18 subfields x 3 sequence types) employed. **Bold** p-values demonstrate significant results.

|  | Left |  |  |  |  |  | Right |  |  |  |  |  |
| --- | --- | --- | --- | --- | --- | --- | --- | --- | --- | --- | --- | --- |
|  | T1 vs T2 |  | T1 vs Slab |  | T2 vs Slab |  | T1 vs T2 |  | T1 vs Slab |  | T2 vs Slab |  |
|  | % | P value | % | P value | % | P value | % | P value | % | P value | % | P value |
| **CA1** | -3.9 | NS | 9.6 | **0.018** | 12.3 | NS | 2.8 | NS | 11.2 | **8.2e-4** | 8.1 | **0.01** |
| **CA2CA3** | -4.1 | NS | 6.0 | NS | 9.5 | NS | 11.6 | **0.0029** | 19.4 | **4.1e-4** | 8.5 | **0.01** |
| **CA4DG** | -<br>11.4 | **0.018** | -<br>13.5 | **4.1e-4** | -2.4 | **0.018** | -5.1 | NS | -9.2 | **8.2e-4** | -4.1 | NS |
| **Subiculum** | 3.0 | NS | 19.0 | **4.1e-4** | 16.0 | NS | 3.7 | NS | 14.7 | **8.2e-4** | 11.1 | **0.020** |
| **SRLM** | -<br>12.6 | **8.2e-4** | 0.2 | NS | 10.6 | **8.2e-4** | -<br>12.8 | **0.0012** | 0.5 | NS | 11.4 | **4.1e-4** |
| **Alveus** | 1.7 | NS | 25.1 | **4.1e-4** | 23.5 | NS | -1.0 | NS | 19.0 | **7.6e-6** | 19.6 | **4.1e-4** |
| **Fimbria** | 7.9 | NS | 17.3 | **4.1e-4** | 9.5 | NS | 15.9 | **0.012** | 24.7 | **0.012** | 10.2 | **0.027** |
| **Fornix** | 0.1 | NS | 31.9 | **4.1e-4** | 30.9 | NS | 3.0 | NS | 36.5 | **7.6e-6** | 33.7 | **4.1e-4** |
| **MB** | 0.0 | NS | 32.7 | **4.1e-4** | 32.2 | NS | -5.1 | NS | 23.7 | **7.6e-6** | 26.9 | **4.1e-4** |

